# Molecular basis of ligand promiscuity, structural mimicry, and atypical dimerization in the chemokine receptors

**DOI:** 10.1101/2024.02.01.578380

**Authors:** Shirsha Saha, Fumiya K. Sano, Saloni Sharma, Manisankar Ganguly, Sayantan Saha, Hiroaki Akasaka, Takaaki Kobayashi, Nashrah Zaidi, Sudha Mishra, Annu Dalal, Samanwita Mohapatra, Manish K. Yadav, Yuzuru Itoh, Rob Leurs, Andy Chevigné, Ramanuj Banerjee, Wataru Shihoya, Osamu Nureki, Arun K. Shukla

## Abstract

Selectivity of natural agonists for their cognate receptors is one of the hallmarks of the members of GPCR family, and it is crucial for the specificity of downstream signal-transduction. However, this selectivity often breaks down in the chemokine receptor subfamily, wherein a high degree of promiscuity is observed with one receptor recognizing multiple chemokines and one chemokine binding to multiple receptors. The molecular determinants of such a striking promiscuity for natural ligands in the chemokine-chemokine receptor system remain mostly elusive and represent an important knowledge gap in our current understanding. Here, we carry out a comprehensive transducer-coupling analysis, testing all known C-X-C chemokines on every C-X-C type chemokine receptor, to generate a global fingerprint of the selectivity and promiscuity encoded within this system. Taking lead from our finding, we determined cryo-EM structures of the most promiscuous receptor, CXCR2, in complex with every interacting chemokine, and deciphered the conserved molecular signatures and distinct binding modalities. While most chemokines position themselves on the receptor as a dimer, CXCL6 exhibits a monomeric binding pose induced by a previously unanticipated reorientation of its carboxyl-terminal α-helix, leading to disruption of the dimer interface. Surprisingly, one of the chemokines, CXCL5, induces a ligand-swapped dimer of CXCR2, the first of its kind observed in class A GPCRs, wherein each protomer of the ligand engages its own receptor without any discernible receptor-receptor interface. These unique observations provide a possible structural mechanism for inherent functional specialization encoded in chemokines despite their convergence to a common receptor. Furthermore, we also determined cryo-EM structures of CXCR3 in complex with G-protein-biased and β-arrestin-biased small molecule agonists that elucidate distinct allosteric modulations in the receptor driving their divergent transducer-coupling bias. Guided by structural analysis and experimental validation, we discover that in contrast to previously held notion, small molecule agonists of CXCR3 display robust agonism at CXCR7, an intrinsically biased, β-arrestin-coupled receptor, making them first-in-class dual agonists for chemokine receptors with exclusive βarr-bias at CXCR7. Taken together, our study provides molecular insights into ligand promiscuity and signaling bias at the chemokine receptors, and also demonstrates a proof of principle that naturally encoded structural mimicry can be recapitulated using synthetic pharmacophores with potential implications for developing novel therapeutics.

## Main

Chemokines are small proteins secreted by immune cells, that play critical roles in a myriad of physiological processes including cellular migration and inflammatory responses by activating chemokine receptors^1,2^. Chemokine receptors belong to the superfamily of G protein-coupled receptors (GPCRs) with primary coupling to Gαi subtype of heterotrimeric G-proteins and β-arrestins (βarrs)^3,4^. They are expressed on a variety of immune cells with wide-ranging contributions to various aspects of our immune response mechanisms, and their aberrant signaling is implicated in multiple disease conditions including cancer^5–7^, allergy^8,9^, psoriasis^10,11^, atherosclerosis^12^, and autoimmune disorders^13,14^. While chemokine receptors exhibit a conserved seven transmembrane architecture characteristic of prototypical GPCRs, their interaction with chemokines does not always follow the exclusive natural agonist selectivity displayed by the majority of GPCRs^15^. This holds true for both C-C and C-X-C type chemokine receptors, as well as for atypical chemokine receptors, such as the Duffy antigen receptor for chemokines, which displays cross-reactivity across C-C and C-X-C chemokines^16–18^. Despite emerging structural insights into chemokine-recognition by chemokine receptors^19–30^, the molecular determinants underlying the inherent ligand promiscuity remain an enigma and represent an important knowledge gap in our current understanding of GPCR activation and signaling paradigm. In this backdrop, we set out to elucidate the molecular mechanism driving ligand promiscuity and selectivity in C-X-C subtype chemokine receptors using a combination of biochemical, pharmacological, and structural approaches.

Considering that the notion of ligand promiscuity at chemokine receptors is based primarily on multiple scattered studies in the literature using different assays and readouts, we first measured the transducer-coupling profile of all known C-X-C chemokines on each of the C-X-C subtype chemokine receptors using G-protein recruitment, βarr2 interaction, and GRK3 recruitment assays in parallel (**Fig. 1a-b**). We observed that CXCR2 exhibits the highest level of promiscuity being activated by seven different chemokines, although the potency and efficacy vary across the ligands (**Fig. 1b-c and Extended Data Fig. 1a-b**). On the other hand, CXCR4 displays the highest degree of selectivity, and is activated only by CXCL12 (**Fig. 1b**). We also observed that despite the high degree of chemokine promiscuity, CXCR2 still maintains some level of selectivity, and fails to exhibit any measurable functional response for several C-X-C chemokines such as CXCL4 and CXCL9-16 (**Fig. 1b**). This is intriguing because the overall structural fold of C-X-C chemokines is highly conserved comprising of three anti-parallel β-strands followed by a carboxyl-terminal α-helix^4^. CXCR2 is expressed on a variety of immune cells including neutrophils, mast cells, monocytes and macrophages, as well as endothelial and epithelial cells^31–34^, and plays an important role in a multitude of cellular and physiological processes such as neutrophil diapedesis, mobilization of neutrophils from the bone marrow to the blood, and neutrophil recruitment in response to microbial infection and tissue injury^35,36^. Thus, it is tempting to speculate that despite chemokine binding promiscuity, there exists some level of functional specialization that fine-tunes context dependent interaction and activation of the receptor. A better understanding of the molecular details of chemokine binding promiscuity and functional specialization may help surmount the inherent challenges in selectively targeting CXCR2 under various disease conditions such as chronic inflammation, cancer progression, psoriasis, atherosclerosis, pulmonary diseases, sepsis, and neuroinflammation^6,10,12,31,37–39^.

**Fig. 1:**
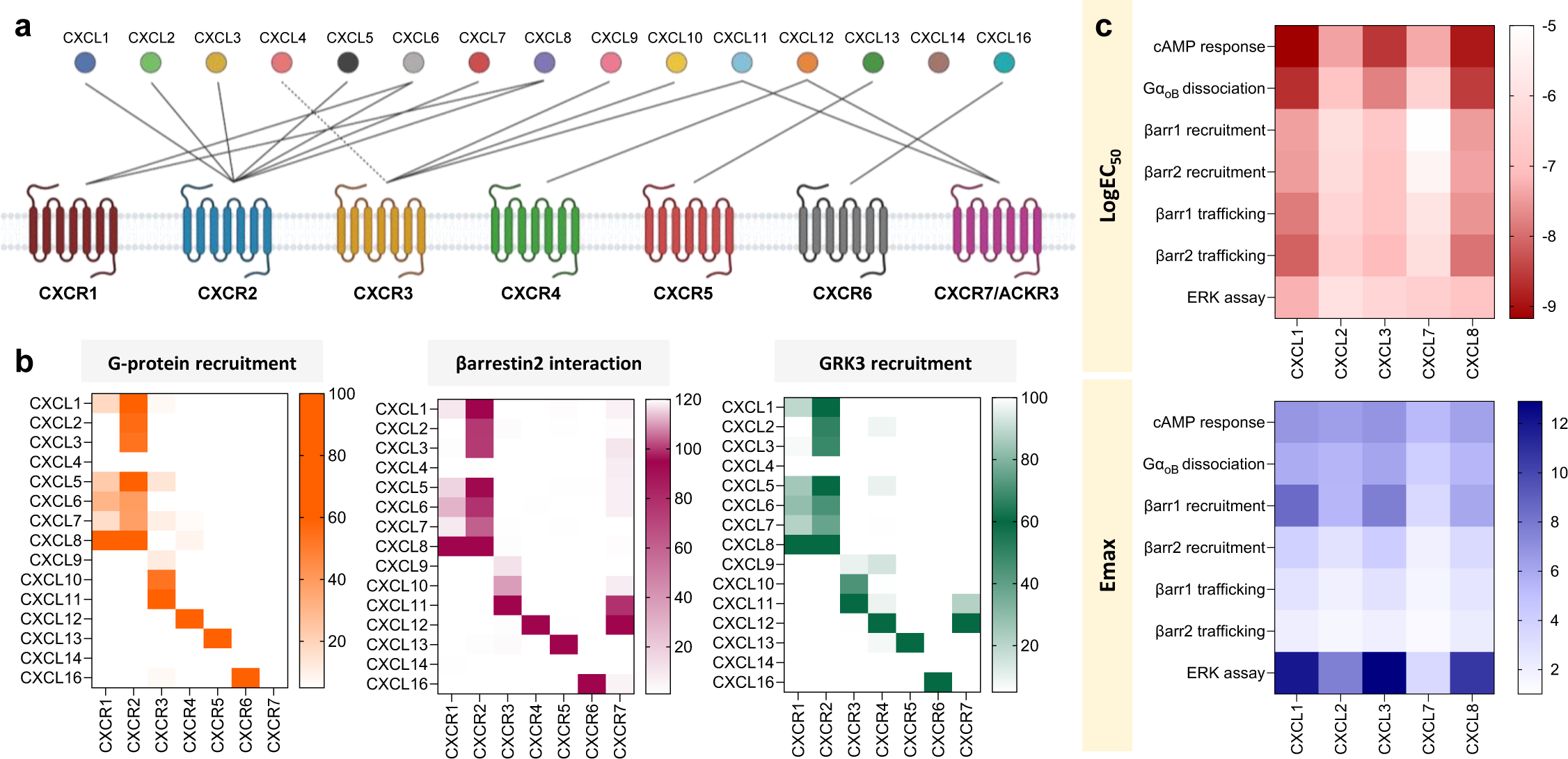
Transducer-coupling profile of all C-X-C chemokines. **a,** Schematic representation of promiscuity and selectivity observed within the C-X-C chemokine receptor family. **b,** Heatmap showing functional selectivity of all C-X-C chemokines on all C-X-C receptors as measured in terms of miniGi, βarr2 and GRK3 recruitment. Data (mean) represents three independent biological replicates normalized with respect to signal observed with most active chemokine agonist, treated as 100%. **c,** Heatmap summarizing the maximal response elicited by CXCR2 downstream to stimulation with different agonists and the respective logEC_50_, in a multitude of assays. Data (mean) represents three-six independent biological replicates, performed in duplicate, and normalized with respect to signal observed at lowest dose, treated either as 100% (for cAMP response and GoB dissociation), or 1 (βarr1/2 recruitment, βarr1/2 trafficking and ERK assay). For cAMP response and GoB dissociation, the decrease observed in luminescence signal was normalized by 10 and plotted.

Taking lead from the chemokine promiscuity fingerprint observed here, we determined the structures of CXCR2 in complex with every interacting chemokine, and heterotrimeric G-protein, using cryo-EM at resolution ranging from 2.8Å to 3.4Å (**Fig. 2a-f, Extended Data Fig. 2a-f, Extended Data Fig. 3-5 and Extended Data Table 1a**). The overall architecture of CXCL-CXCR2-G-protein structures is quite similar and exhibits typical hallmarks of receptor activation such as outward movement of TM6 and rearrangement of conserved motifs (**Extended Data Fig. 6 and 7a**). In addition, the G-protein interaction interface is also similar to what was previously observed for other GPCR-Gαi-protein complexes, and nearly identical across all the CXCR2 structures (**Extended Data Fig. 7c and Extended Data Table 2-7**). Interestingly, we observed two unanticipated features in these structures at the level of chemokine binding modality and receptor dimerization. All the CXCLs except CXCL6 are positioned on the receptor as dimers, wherein one protomer engages the receptor closely while the other protomer points away without making any substantial contact with the receptor (**Fig. 2a-f and 2i, Extended Data Table 2-7**). The dimer interface is conserved across all CXCLs visualized here, and mediated via strong hydrophobic interactions, contributed primarily by the residues from β1 and C-terminal helix of the individual CXCL protomers (**Fig. 2j and Extended Data Fig. 8a-b**). Remarkably, the hydrophobic residues driving these interactions are also conserved in CXCL6 (**Extended Data Fig. 8a-**b). So why does CXCL6 lack a dimeric assembly on the receptor? Structural superimposition of CXCL6 with the other CXCLs reveal that the C-terminal helix in CXCL6 swings outwards by ∼78° from the core domain, and therefore, poses a steric clash with the other protomer in a dimeric assembly (**Fig. 2k**). While chemokines are expected to exist in monomer-dimer equilibrium under physiological conditions^40–42^, it is plausible that their relative dimerization propensity differs from one another, and it may be further fine-tuned upon their interaction with the receptor.

**Fig. 2:**
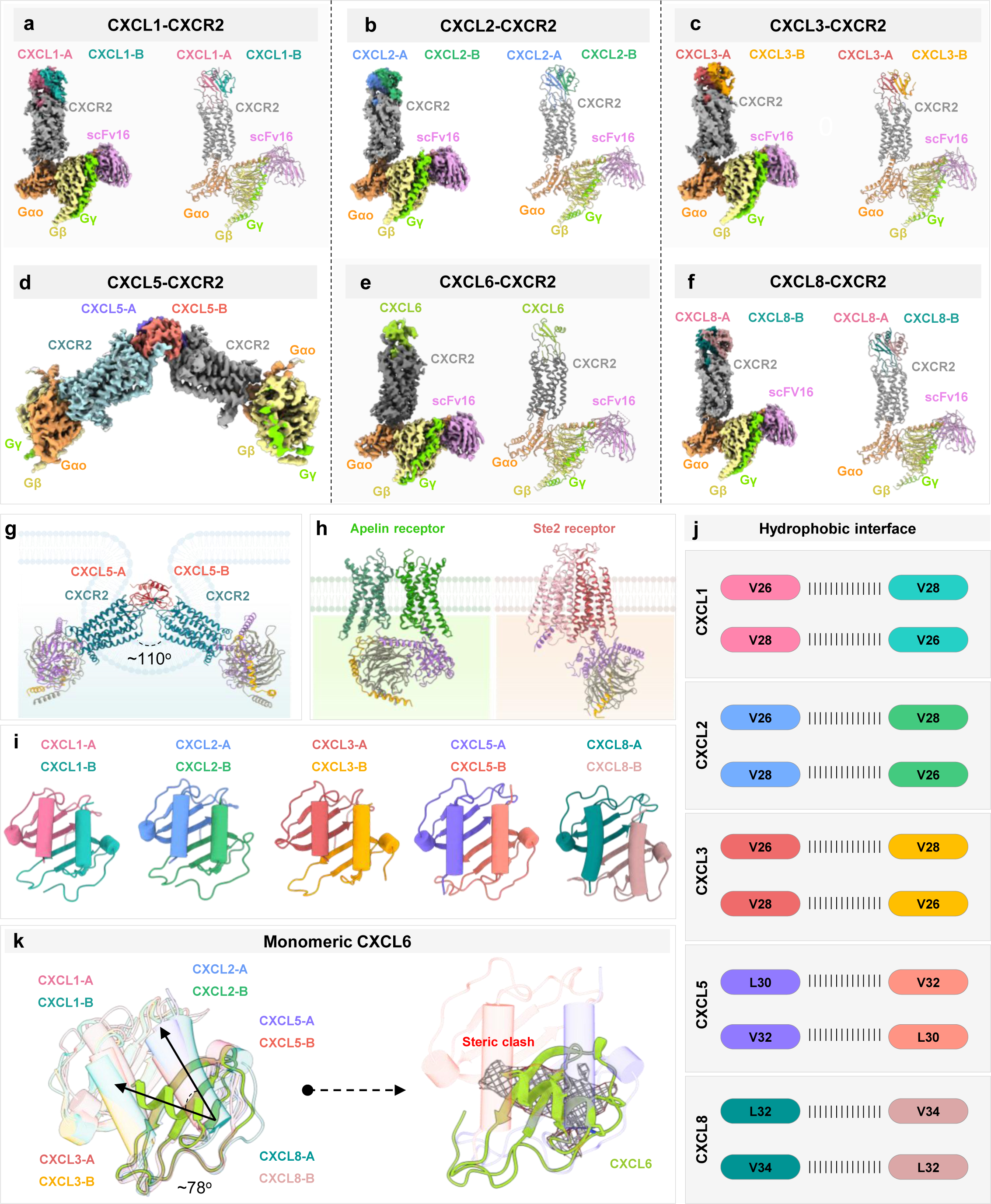
Structures of CXCR2 complexes and ligand conformations. **a-f,** Map and ribbon diagram of the ligand-bound CXCR2-Go complexes (front view) are depicted; **a**, CXCL1-CXCR2-Go: pale violet red: CXCL1-A, light sea green: CXCL1-B, gray: CXCR2, sandy brown: miniGαo, khaki: Gβ1, chartreuse: GL2, plum: scFv16, **b**, CXCL2-CXCR2-Go: cornflower blue: CXCL2-A, medium sea green: CXCL2-B, gray: CXCR2, sandy brown: miniGαo, khaki: Gβ1, chartreuse: GL2, plum: scFv16, **c,** CXCL3-CXCR2-Go: indian red: CXCL3-A, orange: CXCL3-B, gray: CXCR2, sandy brown: miniGαo, khaki: Gβ1, chartreuse: GL2, plum: scFv16. **, d,** CXCL5-CXCR2-Go: medium slate blue: CXCL5-A, salmon: CXCL5-B, gray: CXCR2, sandy brown: miniGαo, khaki: Gβ1, chartreuse: GL2, plum: scFv16, **e,** CXCL6-CXCR2-Go: yellow green: CXCL6, gray: CXCR2, sandy brown: miniGαo, khaki: Gβ1, chartreuse: GL2, plum: scFv16, **f,** CXCL8-CXCR2-Go: teal: CXCL8-A, rosy brown: CXCL8-B, gray: CXCR2, sandy brown: miniGαo, khaki: Gβ1, chartreuse: GL2, plum: scFv16. **g,** Structural representation of dimeric CXCL5-CXCR2 in ribbon form inside an invaginating vesicle. **h**, Comparison of the dimeric states of Apelin receptor (PDB: 7W0L) and Ste2 (PDB: 7AD3). **i,** Structural representations of dimeric C-X-C ligands. **j,** Hydrophobic interactions mediating ligand dimerization. **k**, Comparison of the binding mode of CXCL6 with CXCL1, CXCL2, CXCL3, CXCL5 and CXCL8. The C-terminal helix in CXCL6 shows an outward rotation of ∼78° from the core domain providing an explanation for its monomeric state.

Remarkably, the CXCL5-CXCR2 complex forms a dimer wherein the two protomers of the ligand are arranged in a trans-configuration, with each protomer engaging their own receptor molecule characterized by a large, buried surface area (**Fig. 2d and 2g**). This dimeric architecture displays an angle of approximately 110° between the two receptor molecules, with no direct receptor-receptor contact (**Fig. 2d and 2g**). The overall interaction of CXCL5 with CXCR2 in each protomer is nearly identical including the interaction interface, receptor conformation, and G-protein interaction interface (**Extended Data Table 5**). To confirm that the CXCL5-CXCR2 dimer observed is not a result of the high protein concentration used for cryo-EM analysis, we carried out single particle negative staining of CXCL5-CXCR2 complex, with CXCL8-CXCR2 as a reference, at a significantly lower protein concentration. We observed distinct dimeric classes of CXCL5-CXCR2 samples but not CXCL8-CXCR2, with the latter exhibiting solely monomeric assembly (**Extended Data Fig. 8c**). While class C GPCRs are known to form obligate dimers^43^ (**Extended Data Figure 7f**), so far only one class A GPCR, namely the Apelin receptor has been observed in a dimeric assembly in complex with G-proteins, using cryo-EM^44^ (**Fig. 2h**). The class D fungal GPCR Ste2 has also been visualized recently as a dimer in two different stoichiometries^45,46^ (**Fig. 2h**). However, what is worth noting is that these previously resolved dimers are mediated exclusively by receptor-receptor contact interface unlike the CXCL5-CXCR2 dimer that is mediated only through the ligand interface. Considering the inter-receptor protomer angle and orientation, it is plausible that such a dimeric arrangement represents a receptor internalizing through membrane invagination (**Fig. 2g**) or two interacting receptor protomers from adjacent cells, although the same remains to be experimentally validated in future studies. These two observations i.e. monomeric CXCL6 and CXCL5-induced CXCR2 dimer underscore that despite promiscuous binding to CXCR2, some of the C-X-C chemokines may utilize an additional level of structural specialization to fine-tune their functional outcomes in cellular and physiological context. It is also worth noting that CXCL8 binds to another chemokine receptor CXCR1 as a monomer, and the orientation of ECL2 in CXCR1 has been proposed to possibly clash with the second protomer of CXCL8^21^ (**Extended Data Fig. 8d**). The differential orientation of ECL2 in CXCR2, as compared to CXCR1, permits the binding of dimeric CXCL8, highlighting yet another selectivity level existing within the chemokine system (**Extended Data Fig. 8d**).

The interaction of chemokines with chemokine receptors is conceptualized around a two binding site mechanism, which are referred to as chemokine recognition site 1 and 2 (CRS1 and 2)^47^, respectively (**Fig. 3a**). CRS1, constituted primarily of an interaction of the polar groove within the core domain of the chemokines with the N-terminus of the receptor, is crucial for chemokine recognition^48^ (**Fig. 3c**), while on the other hand, CRS2, formed via the positioning of the N-terminus of chemokines in the orthosteric pocket of the receptors is the key driver of receptor activation and signaling (**Fig. 3d-e and Extended Data Fig. 7d**). Additionally, the conserved Pro38 and Cys39 in the N-terminus of the chemokine receptors, immediately preceding TM1, form the ‘PC motif’ that helps impart a shape complementarity to the N-terminal loop of the chemokines, and this interaction is also referred to as CRS1.5^47^. In the CXCR2 structures, the Cys39^N-term^-Cys286^7^.^25^ disulfide bridge in the receptor packs against the conserved disulfide bridges in the chemokines to facilitate the alignment of the N-terminal loop residues of the receptor with the groove residues of the CXCLs (**Extended Data Fig. 7e**). Furthermore, several hydrogen bonds and ionic contacts help stabilize the flexible N-terminus of CXCR2 within the groove of CXCLs as a part of CRS1 (**Fig. 3c and Extended Data Table 2-7**). Intriguingly, the N-terminus of each of the chemokines is positioned in the orthosteric binding pocket at about the same depth as measured in terms of the distance between the conserved Leu residue in the chemokines and Trp^6.48^ in CXCR2 (**Fig. 3b**). The N-terminus of the chemokines exhibit a shallow binding mode upon penetrating into the orthosteric binding pocket and make extensive contacts within the extracellular vestibule of the TMs, forming the CRS2 (**Fig. 3b and d, Extended Data Fig. 7d and Extended Data Table 2-7**). It is interesting to note that the N-terminus of the receptor bound chemokines undergo a conformational transition from a short and compact hook-shape, in the free-state structures, to a wide and extended “U-shaped” conformation at the base of the orthosteric pocket, with the N-terminal residues extending away from the pocket facilitating the interaction of the N-terminal “hook” with the core domain of CXCLs (**Extended Data Fig. 7b**).

**Fig. 3:**
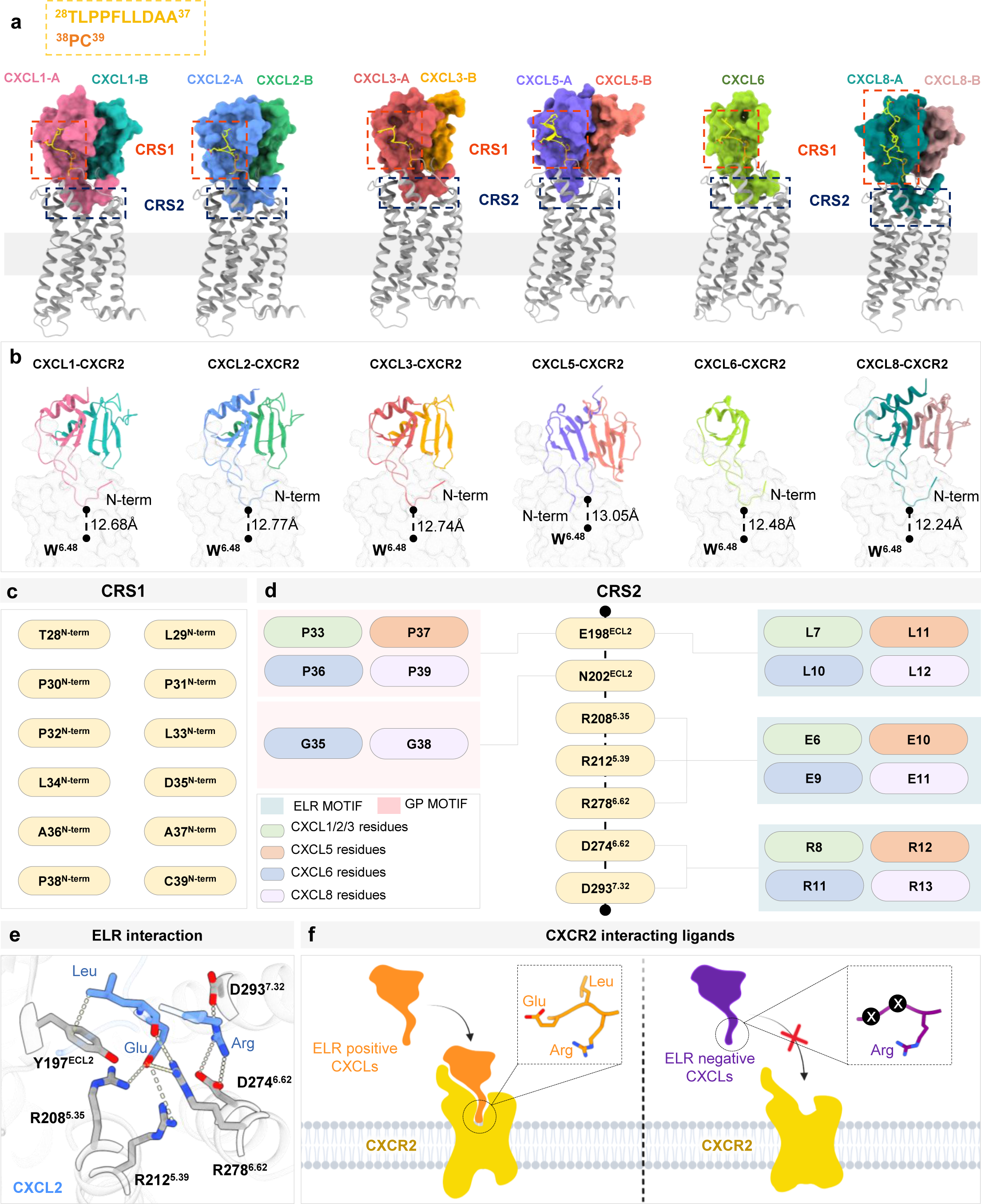
Overall chemokine binding mode in CXCR2. **a**, Representation of the two binding sites engaged by the chemokines on CXCR2. Receptors are shown as foggy ribbon, while chemokines are shown as solid ribbons. CXCR2: gray; CXCL1 protomers: pink, deep cyan; CXCL2 protomers: blue, green; CXCL3 protomers: red, yellow; CXCL5 protomers: purple, salmon; CXCL6: light green; CXCL8 protomers: teal, deep pink. The highly conserved W^6.48^ is highlighted to help infer the depth of insertion of the chemokine N-terminus into the orthosteric pocket of CXCR2. **b,** Binding of individual ligands on CXCR2 and depth with respect to conserved W^6.48^. **c,** Receptor residues in CRS1 which interact with the chemokine. **d**, Residues of CRS2 in CXCR2 interacting with residues of respective chemokine ligands. **e**, Chemokine (CXCL2) ELR residues interacting with CXCR2 residues. **f**, Schematic representation of ELR motif positive ligands interacting with CXCR2.

So, what is the underlying mechanism driving chemokine promiscuity and selectivity at CXCR2? A closer analysis of CXCL-CXCR2 interaction interface provides important insights into this phenomenon. A set of charged residues namely, Arg208^5.35^, Arg212^5.39^, Arg278^6.62^, Asp274^6.58^ and Asp293^7.32^ of CXCR2, hereafter referred to as the “R-D” motif, participate in extensive contacts, through hydrogen bonds and ionic interactions, with the N-terminal ELR motif of CXCLs. Notably, Arg208^5.35^, Arg212^5.39^ and Arg278^6.62^ form polar interactions with the Glu of ELR motif, while Asp274^6.58^ and Asp293^7.32^ interact with the Arg of the ELR motif in every interacting chemokine (**Fig. 3d-e and Extended Data Fig. 7d**). This spatial arrangement and interaction of the “R-D” motif in CXCR2 with ELR motif in CXCL1/2/3/5/6/8 is critical for a common recognition mechanism (**Fig. 3d-e and Extended Data Fig. 7d**). Interestingly, other CXCLs that fail to activate the receptor also lack the ELR motif and thus may not form stable interactions with the receptor amenable to receptor activation (**Fig. 3f and Extended Data Figure 9**). These observations suggest that the spatial positioning of the ELR motif in the angiogenic C-X-C chemokines represents a structural mimicry that facilitates the chemokine promiscuity at CXCR2. It is worth speculating whether other chemokine receptors also follow similar principles of selectivity and promiscuity as observed here for CXCR2.

An intriguing question that remains unanswered is whether the structural promiscuity and mimicry displayed by chemokines can also be recapitulated by small molecule agonists. This is especially important from the perspective of therapeutic targeting of chemokine receptors, which remains challenging and relatively less well explored^49^. CXCR3 is one of the chemokine receptors that is capable of recognizing small molecule scaffolds as agonists, in addition to its natural chemokine agonists^50–52^. CXCR3 is also expressed on a variety of immune cells such as innate lymphocytes, effector T cells, plasmacytoid dendritic cells, subsets of B cells, and within the tumor microenvironment^47,53–56^. Aberrant CXCR3 signaling is implicated in glomerulonephritis, and several inflammatory and neuroinflammatory disorders such as chronic pain, bipolar disorder, rheumatoid arthritis, and spondylitis, making it an important therapeutic target^57–60^. Notably, CXCR3 selectively interacts with only three C-X-C type chemokines i.e. CXCL9-11, which are homeostatic chemokines with angiostatic properties^61^, and in stark contrast to CXCR2, it does not recognize any of the angiogenic ELR motif containing chemokines^16,17^. It is also interesting to note that CXCL11 appears to act as a βarr-biased agonist compared to CXCL9 and CXCL10^62^, and also promotes the formation of non-canonical CXCR3-Gαi-β-arrestin complexes as demonstrated elegantly in cellular context^63^. A splice variant of CXCR3, referred to as CXCR3-B, contains an extended N-terminal domain, and exhibits differential transducer-coupling profile and signaling-bias as compared to the CXCR3-A splice variant^64–67^. Interestingly, a series of small molecule agonists have been described for CXCR3, and transducer-coupling assays have identified VUF10661 as a βarr-biased and VUF11418 as a G-protein-biased agonist (**Fig. 4f**), and they have been reported to exhibit differential responses in terms of chemotaxis and inflammation underscoring their potential therapeutic implications^52^. Therefore, to understand the structural basis of small molecule agonist recognition by CXCR3 and derive insights into their transducer-coupling bias, we determined cryo-EM structures of CXCR3 in apo state, VUF11418, and VUF10661-bound states in complex with heterotrimeric G-proteins (**Fig. 4a-c, Extended Data Fig. 2g-I, Extended Data Fig. 10-11 and Extended Data Table 1b**).

**Fig. 4:**
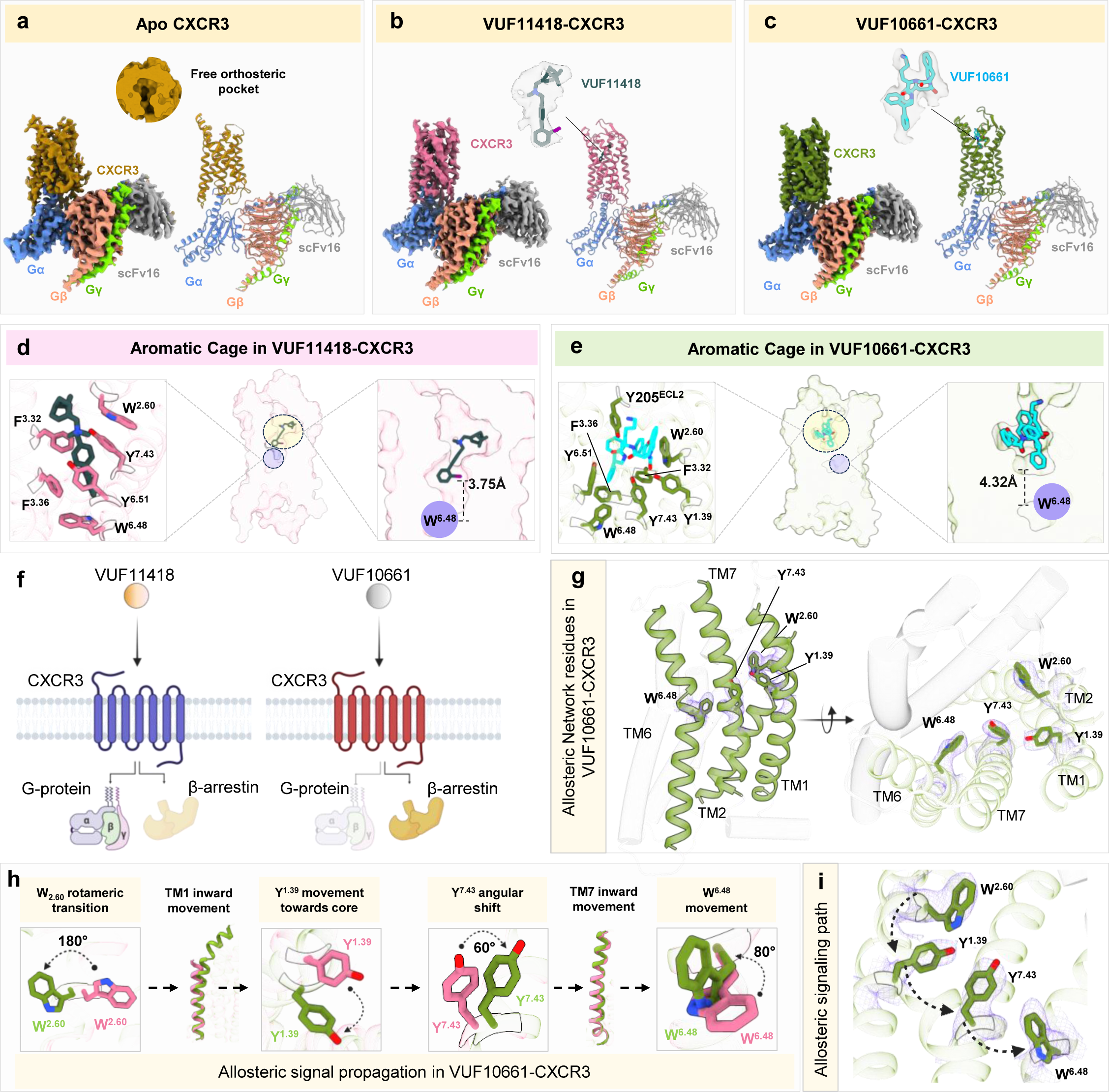
Binding of VUFs on CXCR3 and associated allosteric modulations. **a-c,** Map and ribbon diagram of the apo and ligand-bound CXCR3-Go complexes (front view) and the cryo-EM densities of the ligands (sticks) are depicted as transparent surface representations: **a**, apo-CXCR3-Go: dark goldenrod: CXCR3, cornflower blue: miniGαo, light coral: Gβ1, chartreuse: GL2, grey: scFv16, **b**, VUF11418-CXCR3-Go: pale violet red: CXCR3, cornflower blue: miniGαo, light coral: Gβ1, chartreuse: GL2, grey: scFv16, **c,** VUF10661-CXCR3-Go: olive drab: CXCR3, cornflower blue: miniGαo, light coral: Gβ1, chartreuse: GL2, grey: scFv16. **d-e,** Cross section of the binding pocket of the ligands depicting aromatic cage in CXCR3 and depth with respect to conserved W^6.48^. **f,** Schematic representation of bias exhibited by VUF11418 and VUF10661 upon binding CXCR3. **g,** Key residues in CXCR3 mediating allosteric communication. **h-i,** Allosteric signal propagation in CXCR3 upon binding VUF10661.

The overall structures of CXCR3 are nearly identical to each other in terms of activation dependent conformational changes in the receptor and G-protein binding interface (**Extended Data Fig. 12**), however, there are distinct differences in the agonist-binding mode and local conformations that are linked to downstream transducer-coupling. The ligand binding pocket in CXCR3 is covered by ECL2 at the extracellular surface which adopts a β-hairpin conformation encompassing residues Ser191^ECL2^ to Tyr205^5.53^. Interestingly, VUF10661 adopts an inverted U-shaped binding pose and exhibits a shallower binding mode, as opposed to VUF11418 which penetrates deeper into the orthosteric pocket of the receptor adopting a linear conformation. VUF11418 and VUF10661 occupy a position at a vertical distance of ∼3.8Å and ∼4.3Å, respectively, as measured from the conserved “toggle switch” residue Trp268^6.48^. The ligand binding site in CXCR3 is encapsulated by a cluster of aromatic residues forming an aromatic cage-like structure (**Fig. 4d-e**). A closer analysis of VUF11418- and VUF10661-bound CXCR3 structures reveals a set of networks that are distinct between the two structures. In case of VUF10661, Trp109^2.60^ undergoes a rotameric transition of 180° towards the ligand binding pocket to avoid sterically clashing with Tyr60^1.39^ and to allow the optimal positioning of the ligand. This rotameric shift makes space for the inward movement of the upper portion of TM1 towards the core of the receptor that is relayed further through an angular shift of ∼60° of Tyr308^7.43^ leading to a subsequent inward movement of TM7. These conformational changes allow the rotation of Trp268^6.48^ by 80° towards the ligand binding pocket in case of VUF10661 unlike in VUF11418 (**Fig. 4g-i**). These stark differences in CXCR3 upon binding of VUF10661 vs. VUF11418 hint at an allosteric network connecting the extracellular side of the receptors to the intracellular side through the transmembrane region that directs signaling-bias exhibited by these agonists.

The only other C-X-C type chemokine receptor for which small molecule agonists have been described is CXCR7^29^, which is a βarr-biased receptor with no measurable G-protein coupling but robust βarr recruitment^68^. Taking this into consideration, we compared the key residues in the orthosteric binding pocket of CXCR3 and CXCR7 (**Fig. 5a-b**). Interestingly, we observed a significant conservation of these residues between the two receptors, and it prompted us to probe the reactivity of VUF11418 and VUF10661on CXCR7, and by extension, to the entire panel of CXCRs. Surprisingly, we observed that both VUFs are robust agonists for CXCR7 in βarr recruitment while being silent on G-protein-coupling assays (**Fig. 5c**). Using a previously characterized small molecule agonist of CXCR7, namely VUF11207 as a reference, we further confirmed that VUF11418 and VUF10661 are strong agonists at CXCR7 (**Fig. 5d and Extended Data Figure 1c**). These data suggest that in contrast to previously believed notion, small molecule agonists VUF11418 and VUF10661 are dual agonists of CXCR3 and CXCR7, and therefore, by definition, exclusively biased agonists of CXCR7. It is interesting to note that both CXCR3 and CXCR7 share a common natural chemokine agonist, CXCL11, and our findings with VUF11418 and VUF10661 demonstrate that ligand promiscuity encoded in the natural chemokine agonists can also be recapitulated by engaging only the orthosteric binding pocket by small molecules. At the same time, the exclusive selectivity of VUF11207 for CXCR7 also underscores that selective targeting of the chemokine receptors is also possible, and our structural templates provided here may facilitate efforts in this direction. A direct structural comparison of CXCR3 and CXCR7 structures suggest that the local conformation of the key residues in CXCR3 engaged in interaction with VUF10661, the βarr-biased agonist, align well with the corresponding residues in CXCR7 (**Fig. 5e**). Considering the intrinsic βarr-bias of CXCR7, this observation further supports the contribution of allosteric network and associated local conformational changes in directing transducer-coupling bias at these receptors. These structural correlates also offer a putative template to guide rational design of chemokine receptor targeting entities with signaling bias.

**Fig. 5:**
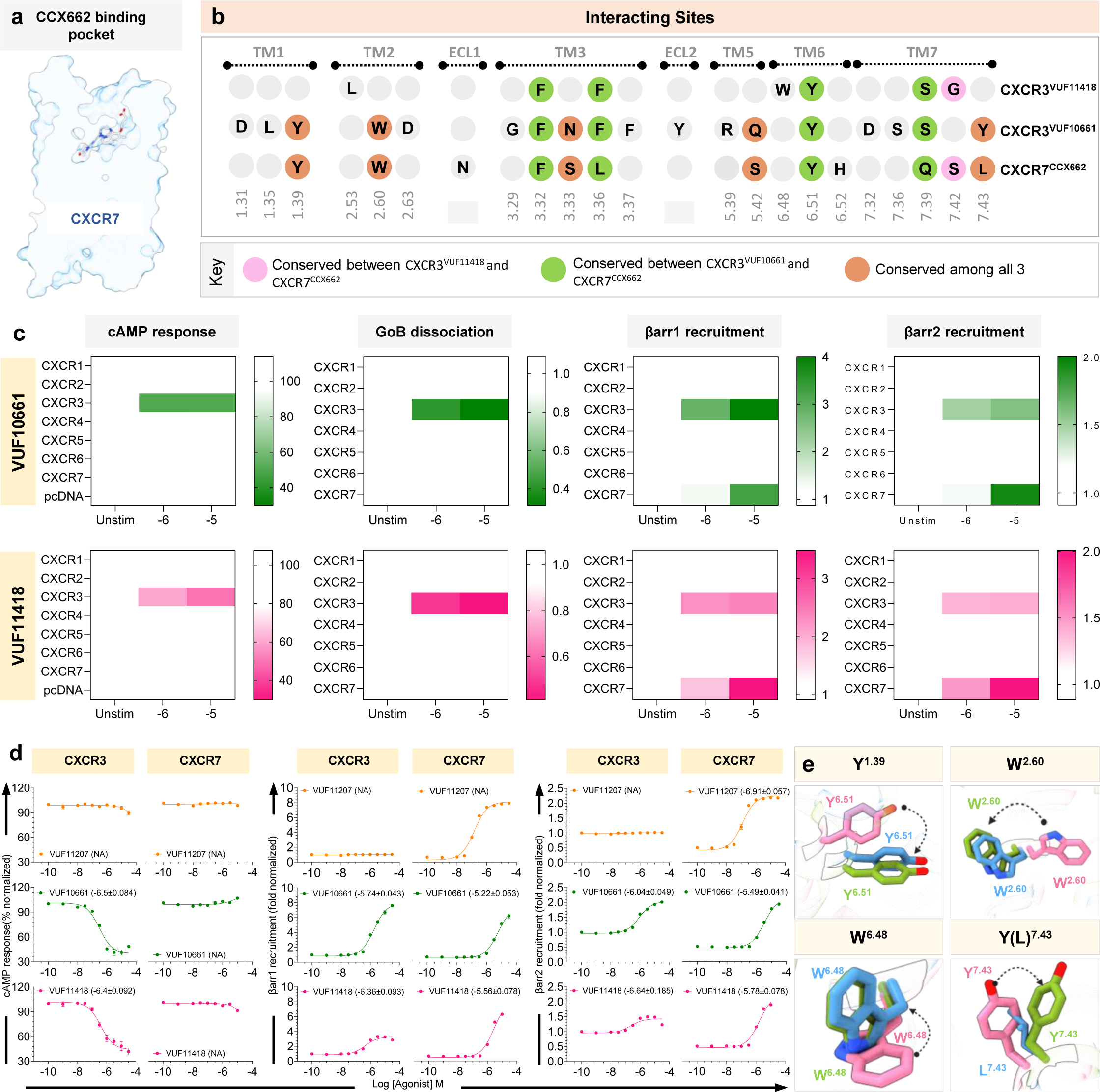
Functional analysis of bias and dual agonism of VUFs. **a,** Cross section of the ligand binding pocket in CCX662 bound CXCR7 (PDB: 7SK9). **b,** Conserved interacting sites in VUF11418-CXCR3, VUF10661-CXCR3 and CCX662-CXCR7. **c,** Heatmap showing VUF11418 and VUF10661 selectivity across all CXCRs in inducing cAMP signaling, GoB dissociation and βarr1/2 recruitment. Data (mean) represents three independent biological replicates, performed in duplicate, and normalized with respect to signal observed in absence of stimulation, treated either as 100% (for cAMP response), or 1 (for GoB dissociation and βarr1/2 recruitment). **d,** VUF11418 and VUF10661 stimulate both CXCR3 and CXCR7, while VUF11207 specifically activates CXCR7, as measured in various assays. Data (mean±SEM) represents three-four independent biological replicates, performed in duplicate, and normalized with respect to signal observed at lowest dose, treated either as 100% (for cAMP response), or 1 (βarr1/2 recruitment). **e,** Residues promoting allosteric communication in VUF10661-CXCR3 (green) exhibit different orientations than those in VUF11418-CXCR3 (pink) and similar rotameric shifts with respect to CCX662-CXCR7 (blue, PDB: 7SK9).

While the current study is focused on C-X-C subtype of chemokine receptors, the C-C chemokine receptors also display a significant level of ligand promiscuity, with some receptors, such as CCR3 binding to more than a dozen different C-C chemokines ^16,17^. It is also striking that some of the C-C chemokine receptors maintain a high degree of ligand selectivity, for example, CCR9, similar to CXCR4 ^16,17^. In addition, some of the chemokine receptors such as the Duffy antigen receptor for chemokine (DARC), also known as the atypical chemokine receptor 1 (ACKR1), even displays cross-reactivity for C-C and C-X-C chemokines^18^. Therefore, the chemokine receptor subfamily represents a rich tapestry for future studies to uncover the fundamental principles that guide naturally encoded ligand-receptor pairing and signaling-bias at multiple levels.

Taken together, our study offers molecular insights into a long-standing dogma of chemokine promiscuity at their receptors, uncovers a non-canonical ligand-swapped dimerization, and a framework for structural mimicry and dual agonism to guide novel ligand discovery at the chemokine receptors with therapeutic potential.

## Methods

### General plasmids, reagents, and cell culture

Most of the molecular biology and general reagents were purchased from Sigma Aldrich unless mentioned otherwise. Dulbecco′s Modified Eagle′s Medium (DMEM), Phosphate buffered saline (PBS), Fetal-Bovine Serum (FBS), Hank’s balanced salt solution (HBSS), Trypsin-EDTA and penicillin-streptomycin solution were purchased from Thermo Fisher Scientific. HEK293T cells (purchased from ATCC, Cat. no: CRL-3216) were maintained in 10cm dishes (Corning, Cat. no: 430167) at 37°C under 5% CO_2_ in Dulbecco’s Modified Eagle’s Medium (Gibco, Cat. no: 12800-017) supplemented with 10% FBS (Gibco, Cat. no: 10270-106), 100U/mL penicillin and 100μg/mL streptomycin (Gibco, Cat. no: 15140-122). *Sf9* cells (purchased from Expression Systems, Cat. no: 94-001LF) were maintained in either ESF921 media (Expression Systems, Cat. no: 96-001-01) or Sf-900^TM^ II SFM serum-free media (Gibco, Cat. no: 10902088). Lauryl Maltose Neopentyl Glycol (L-MNG) was purchased from Anatrace (Cat. no: NG310). The coding regions for CXCR1-7 were sub-cloned in both pcDNA3.1 vector (with an N-terminal FLAG-tag) as well as pCAGGS vector (with an N-terminal FLAG-tag and a C-terminal SmBiT fusion). CXCR2 and CXCR3 were also sub-cloned in pVL1393 vector (with an N-terminal FLAG-tag followed by the N-terminal region of M4 receptor (residues 2-23) which was then used to generate baculovirus encoding the corresponding receptor. The constructs used for NanoBiT-based assays were previously described^69^. All DNA constructs were confirmed by sequencing from Macrogen. VUF11418 and VUF10661 were synthesized and characterized as described previously^50,51^.

### Signal transducers/regulator recruitment assays

Chemokine-induced miniG protein (engineered GTPase domain of Gα subunit)^70^, GRK3^71^ and β-arrestin2^72^ recruitment to chemokine receptors (CXCR1, CXCR2, CXCR3-A, CXCR4, CXCR5, CXCR6 and ACKR3) was monitored using a nanoluciferase complementation-based assay (NanoBiT, Promega)^73,74^. 4×10^6^ HEK293T cells were plated in 10cm dishes and cultured for 24h before transfection with vectors encoding for miniG proteins, human GRK3 or human β-arrestin2 N-terminally fused with LgBiT and the chemokine receptor C-terminally fused with SmBiT. 24h after transfection, cells were harvested, incubated for 15mins at 37°C with coelenterazine H in OptiMEM, and distributed into white 96-well plates (5×10^4^ cells per well). Indicated chemokines (100nM) were then added and the luminescence generated upon nanoluciferase complementation was measured with a Mithras LB940 luminometer (Berthold Technologies) for 20mins. For each receptor, the results are represented as the percentage of the signal monitored with the most active agonist chemokine and presented as mean of three independent experiments (nL=L3).

### Screening all CXCRs with VUF11418/VUF10661

To determine the specificity of VUF11418 and VUF10661, the two ligands were screened against the entire panel of C-X-C receptors in 3 assays: GloSensor Assay (to measure cAMP response), NanoBiT-based G-protein dissociation assay and NanoBiT-based β-arrestin1/2 recruitment assay. HEK293T cells were transfected during splitting. Briefly, the cells were washed with 1X PBS, trypsinized, pooled and resuspended in incomplete media. This was followed by incubation of cells (1.2 million cells for each reaction) with transfection mix and subsequent seeding in 96-well plates at a density of 80,000 cells/well. The transfection mix consisted of either of the following:

- lrl 1μg of N-terminally FLAG-tagged receptor and 1μg of F22 (Promega, Cat. no: E2301) (for GloSensor assay)
- lrl 0.5μg of N-terminally FLAG-tagged receptor, 1μg of GoB tagged with LgBiT at its N-terminus, 1.5μg of Gβ and 1.5μg of GL tagged with SmBiT at its N-terminus (for NanoBiT-based G-protein dissociation assay)
- lrl 1μg of N-terminally FLAG-tagged receptor harboring a C-terminal SmBiT tag and 1μg of either LgBiT-βarr1 or LgBiT-βarr2 (i.e., βarr1/2 harboring an N-terminal LgBiT) (for NanoBiT-based β-arrestin1/2 recruitment assay)

Incomplete media was replaced with complete media after 6-8h. The next day, media was replaced with 100μL assay buffer (For GloSensor assay: 20mM HEPES pH 7.4, 1X Hank’s Balanced Salt Solution/ HBSS and 0.5mg/mL D-luciferin (GoldBio, Cat. no: LUCNA-1G); For NanoBiT assay: 5mM HEPES pH 7.4, 1X HBSS, 0.01% BSA and 10μM coelenterazine (GoldBio, Cat. no: CZ05). The plates were first incubated at 37°C for 1h 30mins followed by an additional 30mins at room temperature.

For GloSensor assay, basal luminescence was measured for 5 cycles using a multiwell plate reader (BMG Labtech). Since we are measuring Gi-mediated decrease in cytosolic cAMP levels, we next added 5μM forskolin to each well, to facilitate an increase in cAMP levels, and measured luminescence for 8 cycles. We then added the different ligands at the indicated final concentration and measured luminescence for 20 cycles.

For NanoBiT-based assays, basal luminescence was recorded for 3 cycles using a multiwell plate reader (BMG Labtech). Ligand was added at the indicated final concentrations and luminescence was recorded for 20 cycles. An average of the luminescence observed for cycles 5-9 was taken. Signal observed was normalized with respect to the luminescence observed at lowest concentration of each ligand, treated as either 100% (for GloSensor assay) or 1 (for NanoBiT assay). Data was plotted and analyzed using GraphPad Prism 10 software.

### GloSensor assay to measure agonist induced decrease in cytosolic cAMP

Agonist induced decrease in cytosolic cAMP levels, as a readout of Gi-mediated second messenger signaling, was measured using GloSensor Assay, as previously described^75–77^. Briefly, HEK293T cells were transfected with 3.5μg of N-terminally FLAG-tagged CXCR2/CXCR3/CXCR7 and 3.5μg of F22 (Promega, Cat. no: E2301). 14-16h post-transfection, the cells were washed with 1X PBS, trypsinized, resuspended in assay buffer (20mM HEPES pH 7.4, 1X Hank’s Balanced Salt Solution/ HBSS and 0.5mg/mL D-luciferin (GoldBio, Cat. no: LUCNA-1G) and seeded in 96-well plates at a density of 100,000 cells/well. This was followed by an incubation of 1h 30mins at 37°C and another 30mins at room temperature. Basal luminescence was then measured for 5 cycles using a multiwell plate reader (BMG Labtech). Since we are measuring Gi-mediated decrease in cytosolic cAMP levels, we next added 5μM forskolin to each well, to facilitate an increase in cAMP levels, and measured luminescence for 8 cycles. We then added the different ligands at the indicated final concentration and measured luminescence for 20 cycles. The signal obtained was normalized with respect to the response obtained at lowest concentration of each ligand, treated as 100%. Data was plotted and analyzed using GraphPad Prism 10 software.

### NanoBiT-based G-protein dissociation assay

Agonist induced G-protein dissociation using a NanoBiT-based assay was measured as previously described^78^. HEK293T cells were transfected with a mixture of 1μg of N-terminally FLAG-tagged CXCR2, 1μg of GoB tagged with LgBiT at its N-terminus, 4μg of Gβ and 4μg of GL tagged with SmBiT at its N-terminus. 14-16h after transfection, the cells were washed with 1X PBS, trypsinized and seeded in 96-well plates at a density of 100,000 cells/well in the presence of assay buffer (5mM HEPES pH 7.4, 1X HBSS, 0.01% BSA and 10μM coelenterazine (GoldBio, Cat. no: CZ05). The plates were first incubated at 37°C for 1h 30mins followed by an additional 30mins at room temperature. Basal luminescence was recorded for 3 cycles using a standard multi-plate reader (Victor X4-Perkin-Elmer). Ligand was added at the indicated final concentrations and luminescence was recorded for 20 cycles. Signal observed was normalized with respect to the luminescence observed at lowest concentration of each ligand, treated as 100%. Data was plotted and analyzed using GraphPad Prism 10 software.

### NanoBiT-based β-arrestin assays

To measure agonist induced β-arrestin1/2 recruitment downstream of CXCR2/CXCR3/CXCR7, we used a previously described NanoBiT-based assay^79,80^. In brief, for measuring β-arrestin1/2 recruitment, HEK293T cells were transfected with 3.5μg of either CXCR2, CXCR3 or CXCR7 (bearing an N-terminal FLAG-tag) and 3.5μg of either LgBiT-βarr1 or LgBiT-βarr2 (i.e., βarr harboring an N-terminal LgBiT). 14-16h after transfection, the cells were washed with 1X PBS, trypsinized and seeded in 96-well plates at a density of 100,000 cells/well in the presence of assay buffer (5mM HEPES pH 7.4, 1X HBSS, 0.01% BSA and 10μM coelenterazine (GoldBio, Cat. no: CZ05). The plates were first incubated at 37°C for 1h 30mins followed by an additional 30mins at room temperature. Basal luminescence was recorded for 3 cycles using a multiwell plate reader (BMG Labtech). Ligand was added at the indicated final concentrations and luminescence was recorded for 20 cycles. An average of the luminescence observed for cycles 5-9 was taken. Signal observed was normalized with respect to the luminescence observed at lowest concentration of each ligand, treated as 1. Data was plotted and analyzed using GraphPad Prism 10 software.

For measuring β-arrestin1/2 trafficking downstream of CXCR2, HEK293T cells were transfected with 3μg of CXCR2 (bearing an N-terminal FLAG-tag), 2μg of N-terminal SmBiT fused β-arrestin1/2 and 5μg of N-terminal LgBiT-fused FYVE.

A NanoBiT-based assay was also used for measuring Ib30 and Ib32 reactivity to β-arrestin1^81^ upon stimulation with different ligands. The transfection mix comprised of 3μg of CXCR2 (bearing an N-terminal FLAG-tag), 2μg of N-terminal SmBiT fused β-arrestin1 and 5μg of N-terminal LgBiT-fused Ib30 or Nb32. The rest of the methodology is the same as described above.

### Measuring ERK signaling using an SRE reporter assay

For measuring ERK signaling downstream to stimulation of CXCR2/CXCR3/CXCR7 with different ligands, we undertook an SRE reporter assay^82^. HEK293T cells were transfected with 3.5μg of N-terminally FLAG-tagged receptor and 3.5μg of an SRE-based luciferase reporter plasmid pGL4.33 (Promega, Cat. no: E1340). 14-16h after transfection, cells were washed with 1X PBS, trypsinized and seeded in 96-well plates at a density of 100,000 cells/well in the presence of complete media. Cells were allowed to settle for 8h, after which complete media was replaced with incomplete DMEM and cells were subjected to starvation overnight. Following this, indicated final concentrations of the various ligands were added and the plates were incubated at 37°C for 6h. Incomplete media was replaced with the assay buffer (20mM HEPES pH 7.4 and 1X HBSS) supplemented with 0.5mg/mL D-luciferin. Luminescence was recorded immediately in a microplate reader (BMG Labtech). Signal observed was normalized with respect to the luminescence observed at lowest concentration of each ligand, treated as 1. Data was plotted and analyzed using GraphPad Prism 10 software.

### Measuring **β**-arrestin recruitment using TANGO assay

To validate that the dual agonism exhibited by VUF10661 and VUF11418 is not an experimental artefact, we measured β-arrestin2 recruitment to CXCR7 using TANGO assay^83^. In brief, HTLA cells were transfected with 7μg of CXCR7 harboring an N-terminal FLAG-tag and a C-terminal TEV protease cleavage site followed by tTA transcription factor. 24h post-transfection, cells were trypsinized and seeded in 96-well plates at a density of 100,000 cells/well in complete DMEM media. After another 24h, complete media was replaced with incomplete media and cells were stimulated with indicated concentration of ligand for an additional 6h at 37°C. Following this, media in the wells was replaced with the assay buffer (20mM HEPES pH 7.4, 1X Hank’s Balanced Salt Solution/ HBSS and 0.5mg/mL D-luciferin (GoldBio, Cat. no: LUCNA-1G). Luminescence was recorded immediately in a microplate reader (BMG Labtech). Signal observed was normalized with respect to the luminescence observed at lowest concentration of each ligand, treated as 1. Data was plotted and analyzed using GraphPad Prism 10 software.

### Receptor surface expression

Receptor surface expression was measured using whole cell ELISA^84^. HEK293T cells expressing FLAG-tagged receptor were seeded in 24-well plates at a density of either 0.1 or 0.2 million cells/well and allowed to adhere overnight. The next day, media was removed from the wells and cells were washed once with 400µL 1X TBS. Cells were fixed by incubating with 300µL of 4% (w/v) paraformaldehyde/PFA for 20mins and excess PFA was removed by washing thrice with 400µL 1X TBS. Wells were blocked with 200µL 1% BSA prepared in 1X TBS for 1h and then incubated with anti-FLAG M2-HRP (1:10,000) (Sigma-Aldrich, Cat. no: A8592) for another 1h. Excess antibody was removed by washing thrice with 400µL 1% BSA. Signal was developed by adding 200µL of tetramethylbenzidine (TMB) (Thermo Fisher Scientific, Cat. no: 34028). Once adequate color developed, the reaction was quenched by transferring 100µL of the solution to a 96-well plate containing 100µL of 1M H_2_SO_4_. Absorbance was recorded at 450 nm using a multimode plate reader (Victor X4-Perkin-Elmer). In order to normalize the response observed across wells, cell density was quantified using Janus Green. Excess TMB solution was removed from the wells and the wells were washed once with 400µL of 1X TBS. Thereafter, the wells were incubated with 200µL of 0.2% (w/v) Janus Green for 15-20mins. Excess stain was removed by washing three times with distilled water and color was developed by adding 800µL of 0.5N HCl to each well. 200µL of this colored solution was transferred to a 96-well plate and absorbance was recorded at 595 nm. Surface expression of the receptor was normalized by taking the ratio of signal observed at 450 nm to signal observed at 595 nm. For all cellular experiments, receptors were expressed at the cell surface at comparable levels (**Extended Data Fig. 13).**

### Purification of chemokines

Coding regions of the various chemokines were cloned in pGEMEX-1 vector with a 6X-His-tag at the N-terminus followed by an enterokinase cleavage site. *E.coli* BL21 (DE3) competent cells were used for over-expression. Transformed cells were inoculated in 50mL TB media containing 100µg/mL ampicillin at 27°C overnight. Primary culture was then inoculated in 1L TB media containing 100µg/mL ampicillin at 27°C until OD_600_ reached 1.5. The culture was then induced with 1mM IPTG and allowed to grow at 20°C for an additional 48h.

For CXCL1/CXCL2/CXCL3/CXCL5/CXCL7/CXCL8/CXCL10, a previously published protocol was followed^85^. Harvested cells were resuspended in lysis buffer (20mM HEPES pH 7.4, 1M NaCl, 10mM Imidazole, 0.3% Triton-X, 1mM PMSF and 5% glycerol) and the resuspension was stirred for 30mins at 4°C. Complete lysis of the cells was achieved by ultrasonication for 20mins. This was followed by centrifugation at 18,000 rpm at 4°C for 30mins to remove the cell debris. Protein was enriched on Ni-NTA beads, and excess unbound/non-specific protein was removed by washing with wash buffer (20mM HEPES pH 7.4, 1M NaCl, 40mM Imidazole and 5% glycerol). Protein was eluted with elution buffer (20mM HEPES pH 7.4, 100mM NaCl, 500mM Imidazole and 5% glycerol) and the eluate was dialyzed against enterokinase digestion buffer (20mM Tris-Cl pH 7.5, 150mM NaCl and 2.5% Glycerol) overnight at 4°C. Precipitated protein was removed by centrifugation at 5000 rpm at 4°C for 10mins. Digestion was set up to remove the 6X-His-tag by incubating with either homemade or store bought (NEB, Cat. no: P8070L) enterokinase in the presence of 10mM CaCl_2_ at 22°C for 16h. Cleaved protein was then loaded onto the Resource S Cation Exchange Chromatography column (Cytiva Life Sciences, Cat. no: 17118001) (Loading buffer: 50mM MES pH 5.5, 50mM NaCl). Before loading, salt was diluted 3x using 100mM MES buffer pH 5.5. Gradient elution was taken by generating a linear gradient of NaCl (100mM-1M) over 16 column volumes. Peak fractions were pooled on the basis of SDS-PAGE and then dialyzed against PD-10 buffer (20mM HEPES pH 7.4, 150mM NaCl) overnight at 4°C. Protein was flash frozen and stored at -80°C in the presence of 10% glycerol.

For CXCL5, following enterokinase cleavage the protein was concentrated and loaded onto HiLoad Superdex 16/600 200 PG column (Cytiva Life sciences, Cat. no: 17517501). Fractions corresponding to cleaved CXCL5 were pooled, flash-frozen and stored at -80°C in the presence of 10% glycerol.

For purifying CXCL6, every 10g of pellet was resuspended in 50mL of Buffer A (50mM Tris-HCl pH 8.0, 6M guanidinium HCl pH 8.0 and 200mM NaCl). The cells were allowed to solubilize for a period of 1h at 4°C and then lysed by sonication. The cell lysate was then isolated via centrifugation at 25,000 rpm for 40mins and then applied to a Ni-NTA column. The beads were then washed with 2 CVs of Buffer B (6M guanidinium HCl pH 8.0 and 200mM NaCl) and eluted with Buffer C (20mM Tris-HCl pH 8.0, 200 mM NaCl and 500 mM imidazole). The eluted protein was then incubated with 20mM DTT for an hour and was then diluted dropwise in Buffer D (0.55M L-arginine hydrochloride, 20mM Tris-HCl, 200mM NaCl, 1mM EDTA, 1 mM reduced glutathione and 0.1mM oxidised glutathione pH 8.0) and incubated for 48h at 4°C. The protein solution was then concentrated with Vivaspin 10kDa MWCO concentrator (Cytiva Life sciences, Cat. no: 28932360) and dialysed against 20mM Tris-HCl pH 8.0, 200mM NaCl. The amount of protein was estimated by running SDS-PAGE and then digestion reaction was set up with homemade enterokinase, supplemented with 10mM CaCl_2_. The enterokinase digested CXCL6 was then concentrated with Vivaspin MWCO 3kDa (Cytiva Life sciences, Cat. no: 28932293) and then injected into HiLoad Superdex 16/600 200 pg column (Cytiva Life sciences, Cat. no: 17517501). Fractions corresponding to the protein were pooled, flash-frozen and stored at -80°C with 10% glycerol.

### Expression and Purification of Enterokinase

A DNA construct of bovine enteropeptidase catalytic light chain with N terminal-Trx tag followed by Thrombin cut site and a self-cleavable enterokinase site was cloned in pET-32a (+) vector. 6X-His-tag was present at the C-terminal end of the protein and a mutation was introduced in the 112^th^ residue to change it from C to S. The DNA was transformed in *E. coli* SHuffle strain and a single isolated colony from the transformed plate was inoculated in 50mL of LB media and allowed to grow overnight at 30°C. The primary culture was then transferred to 0.5L of TB media followed by induction with 70μM of IPTG at an optical density of 0.7 and allowed to grow for 16h at 16°C. Culture flasks were supplemented with a final concentration of 100μg/mL of freshly prepared ampicillin. The cells were then harvested by centrifugation after 18h and resuspended in 50mL of resuspension buffer (20mM Tris-HCl pH 7.5, 10mM EDTA, 1% triton-X-100 and 2mM CaCl_2_) and were allowed to solubilise for a period of 30mins at 4°C. Cells were lysed by sonication and the supernatant was separated by centrifugation for 30mins at 20,000 rpm at 4°C. The pellet obtained was then dissolved in 10mL of 0.1M Tris-HCl pH 8.6, 1mM EDTA, 20mM DTT and 6 M guanidinium HCl. The insoluble fractions were separated by centrifugation at 25,000 rpm for 20mins at 4°C. The supernatant was collected and put up for dialysis against 3M guanidinium HCl pH 2.5 at room temperature. After dialysis the solution was mixed with 10mL of oxidation buffer (50mM Tris-HCl pH 8.3, 6M guanidinium-HCl, 0.1M oxidised glutathione) and then again dialysed against 3M guanidinium HCl pH 8.0. For initiating the refolding process, the dialysed protein solution was then dropwise diluted into 600mL of 0.7M L-arginine hydrochloride pH 8.6, 2mM Reduced glutathione and 1 mM EDTA and then incubated for 75h at 4°C. The protein was then subsequently dialysed against 0.1M Tris-HCl and 10 mM CaCl_2_ and loaded onto Ni-NTA column, washed with 10mM Tris-HCl, 500mM NaCl and eluted with 500mM Imidazole containing elution buffer. The elution was then dialysed against and finally stored in 0.1M Tris-HCl pH 8.0, 500mM NaCl and 50% glycerol at -20°C.

### Purification of CXCR2 and CXCR3

Full length recombinant CXCR2/CXCR3 was isolated from *Spodoptera frugiperda (Sf9)* insect cells following a previously published protocol^18,69,86^. *Sf9* cells were harvested 72h post-infection with CXCR2/CXCR3 expressing baculovirus. This was followed by homogenisation of the cells initially in hypotonic buffer (20mM HEPES pH 7.4, 20mM KCl, 10mM MgCl₂, 1mM PMSF, 2mM benzamidine) and subsequently in hypertonic buffer (20mM HEPES pH 7.4, 20mM KCl, 10mM MgCl2, 1M NaCl, 1mM PMSF, 2mM benzamidine). Cells were then subjected to solubilization by incubating in lysis buffer (20mM HEPES pH 7.4, 450mM NaCl, 1mM PMSF, 2mM benzamidine, 0.1% cholesteryl hemisuccinate, 2mM iodoacetamide and 0.5% L-MNG (Anatrace, Cat. no: NG310) for 2h at 4°C. Next, the lysate was diluted in 2 times volume of dilution buffer (20mM HEPES pH 7.4, 8mM CaCl₂, 1mM PMSF, and 2mM benzamidine) to reduce the salt concentration to 150mM NaCl. Debris was removed by centrifuging the lysate at 20,000 rpm for 30mins. The supernatant was filtered and loaded onto pre-equilibrated M1-FLAG beads. The column was then washed alternatively with LSB/ Low Salt Buffer (20mM HEPES pH 7.4, 150mM NaCl, 2mM CaCl₂, 0.01% cholesteryl hemisuccinate, 0.01% L-MNG) and HSB/ High Salt Buffer (20mM HEPES pH 7.4, 350mM NaCl, 2mM CaCl₂, 0.01% L-MNG). Protein was eluted in the presence of 2mM EDTA and 250μg/mL FLAG. To prevent receptor aggregation, free cysteines were blocked by incubating with 2mM iodoacetamide. Excess free iodoacetamide was quenched by incubating with 2mM L-cysteine.

Apo purified CXCR2 was incubated with either 1.5X molar excess (for CXCL1, CXCL2, CXCL3, CXCL5, CXCL8) or 3X molar excess (for CXCL6) of chemokine for 1h at room temperature. For CXCR3, ligand (either 100nM CXCL10 or 1µM VUF11418 or 1µM VUF10661) was kept in all the buffers during purification. Ligand bound receptor was stored in the presence of 10% glycerol at -80°C till further use.

### Purification of G-proteins

MiniGαo was purified from *E. coli* BL21 (DE3) cells according to a previously published protocol^69,86^. A starter culture was grown for 6-8h at 37°C in LB media, followed by an overnight primary culture at 30°C in the presence of 0.2% glucose supplementation. Secondary culture was grown in TB/ Terrific Broth media and induced at an OD_600_ of 0.8 with 50µM IPTG. Following induction, cells were cultured for an additional 18-20h at 25°C. Cells thus obtained were lysed by sonication in lysis buffer (40mM HEPES pH 7.4, 100mM NaCl, 10% Glycerol, 10mM Imidazole, 5mM MgCl_2_, 1mM PMSF, 2mM benzamidine, 1mg/mL lysozyme, 50µM GDP and 100µM DTT). Cell debris was removed by centrifuging at 20,000 rpm for 30 mins and the filtered supernatant was enriched on Ni-NTA beads. Excess unbound protein was removed by washing with wash buffer (20mM HEPES pH 7.4, 500mM NaCl, 40mM Imidazole, 10% Glycerol, 50μM GDP and 1mM MgCl_2_) and bound protein was eluted in 20mM HEPES pH 7.4, 100mM NaCl, 10% Glycerol and 500mM Imidazole. 6X-His-tag was removed by treating with TEV protease overnight (TEV:protein, 1:20) at room temperature and cleaved untagged protein was isolated by size exclusion chromatography using HiLoad Superdex 200 PG 16/600 column (Cytiva, Cat. no: 17517501). Fractions corresponding to our protein of interest were pooled, quantified and stored in the presence of 10% glycerol at -80°C till further use.

Gβ1γ2 was purified from *Sf9* insect cells as previously described^69,86^. Gβ1 and Gγ2 were co-expressed in *Sf9* insect cells using the baculovirus expression system, with Gβ1 containing an N-terminal His tag. 72h post infection, cells were harvested and lysed by sequentially douncing first in lysis buffer (20mM Tris-Cl pH 8.0, 300mM NaCl, 10% Glycerol, 1mM PMSF, 2mM benzamidine and 1mM MgCl_2_) and then in solubilization buffer (20mM Tris-Cl pH 8.0, 300mM NaCl, 10% Glycerol, 1% DDM, 5mM β-ME, 10mM Imidazole, 1 mM PMSF and 2mM benzamidine). Solubilization was allowed to proceed for 2h at 4°C, which was followed by centrifugation at 20,000 rpm for 30mins to clear cellular debris. The supernatant was filtered and loaded onto pre-equilibrated Ni-NTA beads. Unbound protein was removed by washing extensively with wash buffer (20mM Tris-Cl pH 8.0, 300mM NaCl, 30mM Imidazole, 10% glycerol, 5mM β-ME and 0.02% DDM (Anatrace, Cat. no: D310A) and eluted with 20mM Tris-Cl pH 8.0, 300mM Imidazole and 0.01% L-MNG. Eluted protein was quantified and stored in the presence of 10% glycerol at -80°C till further use.

### Purification of scFv16

Gene encoding scFv16 was cloned in pET-42a (+) vector with an in-frame N-terminal 10X-His-MBP tag followed by a TEV cleavage site and expressed in *E. coli* Rosetta (DE3) strain, following a previously published protocol^69,86,87^. Overnight primary culture was transferred to 1L 2xYT media supplemented with 0.5% glucose and 5mM MgSO_4_. The culture was then induced at an OD_600_ of 0.9 with 250μM isopropyl-β-D thiogalactopyranoside (IPTG) and allowed to grow for 16–18h at 18°C. Cells were resuspended in 20mM HEPES pH 7.4, 200mM NaCl, 2mM Benzamidine, and 1mM PMSF and incubated at 4°C for 40mins with constant stirring. Cells were disrupted by ultrasonication, and cell debris was removed by centrifugation at 18,000 rpm for 40mins at 4°C. Protein was enriched on Ni-NTA resins, and beads were washed extensively with 20mM HEPES pH 7.4, 200mM NaCl, 50mM Imidazole. Bound protein was eluted with 300mM Imidazole in 20mM HEPES pH 7.4, 200mM NaCl. Subsequently, Ni-NTA elute was enriched on amylose resin (NEB, Cat. no: E8021L) and washed with 20mM HEPES pH 7.4, 200mM NaCl to remove non-specific proteins. Protein was then eluted with 10mM maltose prepared in 20mM HEPES pH 7.4, 200mM NaCl, and the His-MBP tag was removed by overnight treatment with TEV protease (TEV protease:Protein 1:20). Tag-free scFv16 was recovered by passing TEV-cleaved protein through Ni-NTA resin. Eluted protein was concentrated with Vivaspin 10kDa MWCO concentrator (Cytiva Life sciences, Cat. no: 28932360) and cleaned by size exclusion chromatography on HiLoad Superdex 16/600 200 PG column (Cytiva Life sciences, Cat. no: 17517501). Fractions corresponding to scFv16 were pooled, flash-frozen and stored at - 80°C in presence of 10% glycerol.

### Reconstituting chemokine/synthetic ligand-chemokine receptor-G protein complexes

Purified chemokine-receptor complex was incubated with 1.2-fold molar excess of Gαo, Gβ1γ2, and scFv16, in the presence of 5mM CaCl_2_ and 25mU/mL apyrase (NEB, Cat. no: M0398S), for 2h at room temperature. The mixture was then concentrated using a 100 MWCO concentrator (Cytiva, Cat. no: GE28-9323-19) and injected into Superdex200 Increase 10/300 GL SEC column to separate the receptor-G-protein complex from the free components. Peak fractions were analyzed by running an SDS-PAGE. Fractions containing the complex were pooled and concentrated to roughly 12-18mg/mL using the same concentrator and stored at -80°C until further use.

### Negative stain electron microscopy

Prior to grid freezing for high-resolution cryo-EM data collection, conventional uranyl-formate negative staining was used to assess sample homogeneity^79^ ^18^. In brief, a formvar/carbon-coated 300 mesh copper grid (PELCO, Ted Pella) was dispensed with 3.5µl of the sample, incubated for 1 minute, and then blotted off using Whatman No. 1 filter paper. The grid containing the attached sample was then touched onto a first drop of freshly prepared 0.75% uranyl formate stain, and immediately blotted off using filter paper. To improve staining efficiency, the grid was then placed on a second drop of uranyl formate and moved in a rotating fashion for 30 seconds. Before imaging and data collection, the excess stain was blotted off and allowed to air dry. A FEI Tecnai G2 12 Twin TEM (LaB6) operating at 120kV and outfitted with a Gatan 4k x 4k CCD camera at 30,000x magnification was used for imaging and data collection. For further analysis, the acquired micrographs were imported into Relion 3.1.2^88–90^. About 10,000 particles were automatically selected using the Gaussian blob picker, extracted with a box-size of 280 pix, and then submitted to reference-free 2D classification to obtain the final 2D class averages.

### Cryo-EM grid preparation and data collection

3.0µl of the purified CXCR3-Go and CXCR2-Go complexes were dispensed onto glow discharged Quantifoil holey carbon grids (R1.2/1.3, Au, 300 mesh) at a concentration of approximately 13.2 mg/ml (VUF10661-CXCR3-Go), 18.5 mg/ml (VUF11418-CXCR3-Go), 14.7 mg/ml (apo-CXCR3-Go), 15.0 mg/ml (CXCL1-CXCR2-Go), 16.7 mg/ml (CXCL2-CXCR2-Go), 12.1 mg/ml (CXCL3-CXCR2-Go), 16.6 mg/ml (CXCL5-CXCR2-Go), 18.4 mg/ml (CXCL6-CXCR2-Go), and 23.4 mg/ml (CXCL8-CXCR2-Go). The grids were blotted for 4 seconds at 4°C and 100% humidity with a blot force of 10 using a Vitrobot Mark IV (Thermo Fischer Scientific) and immediately plunge frozen in liquid ethane (-181°C).

Data collection of all samples were performed on a Titan Krios G3i (Thermo Fisher Scientific) operating at an accelerating voltage of 300kV equipped with a Gatan K3 direct electron detector and BioQuantum K3 imaging filter. Movie stacks were acquired in counting mode at a pixel size of 0.83 Å/pix and a dosage rate of approximately 15.6 e^-^/Å^2^/s using EPU software over a defocus range of -0.8 to -1.6μm. Each movie was fractionated into 48 frames with a total dose of 50.1 e^-^/Å^2^ that was obtained throughout the 2.3 s exposure period. In total, 3165, 3030, and 3125 movie stacks were collected for VUF10661-CXCR3, VUF11418-CXCR3 and Apo-CXCR3 samples respectively, while 8,273, 3,555, 2,108, 3,752, 4,722, and 4,509 movie stacks were acquired for CXCL1-CXCR2-Go, CXCL2-CXCR2-Go, CXCL3-CXCR2-Go, CXCL5-CXCR2-Go, CXCL6-CXCR2-Go, and CXCL8-CXCR2-Go respectively.

### Cryo-EM data processing

Movie stacks were aligned (4×4 patches) and dose-weighted using RELION’s implementation of the MotionCor2 algorithm^90^. The motion-corrected micrographs were imported into cryoSPARC v4.4^91^, and CTF parameters were estimated with Patch CTF (multi).

For the VUF10661-CXCR3-Go dataset, 1,384,864 autopicked particle projections were extracted using a box size of 280 pix (fourier cropped to 70 pix) and then subjected to 2D classification for cleaning. 363,327 particle projections corresponding to 2D class averages with evident secondary features were selected, re-extracted with a box size of 280 pix (fourier cropped to 180 pix), and subjected to heterogeneous refinement into 3 classes. The particles curated through several rounds of heterogeneous refinement were exported to RELION v4.0. Subsequently, further curation was performed, with a focus on the receptor region, followed by Bayesian polishing with a box size of 300 pix (fourier cropped to 240 pix). The 116,462 resulting particles were imported into cryoSPARC and subjected to non-uniform refinement with estimating CTF parameters, yielding a reconstruction with a nominal resolution of 3.03 Å at a fourier shell correlation of 0.143. In order to improve the resolution and features corresponding to the receptor, local refinement was performed with a mask on the receptor, yielding a reconstruction with a nominal resolution of 3.16 Å.

For the VUF11418-CXCR3-Go dataset, 1,527,953 particles were autopicked from 3,030 motion-corrected micrographs using the template-picker subprogram within cryoSPARC. Picked particles were extracted with a box size of 280 pix and fourier cropped to 70 pix, and subjected to 2D classification and heterogeneous refinement to remove ice contamination and dissociated particles. The resulting 360,223 particles were re-extracted with a box size of 280 pix (fourier cropped to 180 pix) and subjected to heterogeneous refinement into 3 classes. The 182,526 resulting particles were exported to RELION v4.0. Subsequently, further curation was performed, with a focus on the receptor region, followed by Bayesian polishing with a box size of 300 pix (fourier cropped to 240 pix). The 150,213 resulting particles were imported into cryoSPARC and subjected to non-uniform refinement with estimating CTF parameters, yielding a reconstruction with a nominal resolution of 3.07 Å at a fourier shell correlation of 0.143. Local refinement of the receptor region with a mask improved the density derived from receptor, yielding a reconstruction with a nominal resolution of 3.53 Å.

For the Apo-CXCR3-Go dataset, template-picker was used to automatically pick particles from 3,125 motion-corrected micrographs. The 1,633,141 picked particles were extracted with a box size of 280 pix (fourier cropped to 70 pix) and subjected to 2D classification and heterogeneous refinement to remove the contaminations and dissociated particles. The resulting 298,771 particles were re-extracted with a box size of 280 pix and fourier cropped to 180 pix followed by heterogeneous refinement. The 173,083 resulting particles were exported to RELION v4.0. Subsequently, further curation was performed, with a focus on the receptor region, followed by Bayesian polishing with a box size of 300 pix (fourier cropped to 240 pix). The 41,722 resulting particles were imported into cryoSPARC and subjected to non-uniform refinement with estimating CTF parameters, yielding a reconstruction with a nominal resolution of 3.30 Å at a fourier shell correlation of 0.143. To improve the resolution and features corresponding to the receptor, local refinement was performed with a mask on the receptor, yielding a reconstruction with a nominal resolution of 3.68 Å.

All datasets of the CXCR2-Go complexes were processed following a similar pipeline as that of CXCR3. Briefly, raw movies were aligned with MotionCor2 in RELION 4.0, imported into cryoSPARC v4.4 and subjected to CTF estimation using Patch CTF (multi). For the CXCL1-CXCR2-Go dataset, 4,437,786 autopicked particles (template based) were extracted using a box size of 280 pix (fourier cropped to 70 pix) and then cleaned using reference-free 2D classification and heterogeneous refinement to remove ice contamination and distorted particles. 317,394 particles were re-extracted with a box size of 280 pix (fourier cropped to 180 pix), and subjected to heterogeneous refinement into three 3 classes. 115,169 particles that were curated via many rounds of heterogeneous refinement were imported into RELION v4.0. Following Bayesian polishing with a box size of 300 pix (fourier cropped to 240 pix), the polished particles were imported into cryoSPARC and subjected to CTF refinement and NU refinement, providing a reconstruction with a nominal resolution of 3.07 Å at fourier shell correlation of 0.143. To further improve the density of the receptor region, local refinement was performed using the receptor-focused mask, providing a reconstruction with a nominal resolution of 3.48 Å.

For the CXCL2-CXCR2-Go dataset, 1,927,680 template based autopicked particles were extracted using a box size of 280 pix (fourier cropped to 70 pix) and then cleaned using heterogeneous refinement to remove ice contamination and distorted particles. 623,954 particle s were re-extracted with a box size of 280 pix (fourier cropped to 180 pix), and subjected to heterogeneous refinement into three 3 classes. 285,884 particles corresponding to the best 3D class were imported into RELION v4.0. Following Bayesian polishing with a box size of 300 pix (fourier cropped to 240 pix), the resultant polished particles were imported into cryoSPARC and subjected to CTF refinement and NU refinement (in cryoSPARC), providing a reconstruction with a nominal resolution of 2.8 Å at 0.143 fourier shell cut-off. To further improve the density of the receptor region, local refinement was performed using the receptor-focused mask, providing a reconstruction with a nominal resolution of 3.09 Å.

For the CXCL3-CXCR2-Go dataset, 1,133,660 template picked particles were extracted using a box size of 280 pix (fourier cropped to 70 pix) and then cleaned using heterogeneous refinement to remove ice contamination and distorted particles. 307,000 particles were re-extracted with a box size of 280 pix (fourier cropped to 180 pix), and subjected to heterogeneous refinement into three 3 classes. RELION v4.0 was used to import the particles curated via many rounds of heterogeneous refinement, and subjected to 3D classification without alignment followed by Bayesian polishing with a box size of 300 pix (fourier cropped to 240 pix). 46,110 particles that resulted were imported into cryoSPARC and subjected to non-uniform refinement using estimated CTF values, providing a reconstruction with a nominal resolution of 3.38 Å at a fourier shell correlation of 0.143. Local refinement using a mask on the receptor was performed to improve the features corresponding to the receptor, providing a reconstruction with a nominal resolution of 3.65 Å.

For the CXCL5-CXCR2-Go dataset, 2,047,293 particles were automatically picked using the template picker subprogram, extracted with a box size of 560 pix (fourier cropped to 140 pix) and subjected to several rounds of 2D classification. Following re-extraction with a box size of 560 pix (fourier cropped to 320 pix), and heterogeneous refinement with a C2 symmetry constraint were performed to remove fuzzy particles, yielding a total of 131,780 particles. The clean particle stack was imported into RELION v4.0 subjected to Bayesian polishing with a box size of 560 pix (fourier cropped to 440 pix), particles were imported back into cryoSPARC. Imported particles were subjected to CTF refinement and NU refinement with a C2 symmetry constraint to produce a map with a global indicated resolution of 3.32 Å at fourier shell correlation of 0.143. Local refinement with a mask on the receptor with a C2 symmetry constraint was performed to improve the interpretability of the map, yielding a reconstruction with a global resolution of 3.06 Å.

For the CXCL6-CXCR2-Go dataset, 1,609,421 particles were picked and extracted with a box size of 280 pix (fourier cropped to 70 pix), and subjected to several rounds of heterogeneous refinement to eliminate carbon edges and ice contaminations in cryoSPARC. Following re-extraction with a box size of 280 pix (fourier cropped to 180 pix), and subjected to heterogeneous refinement into three 3 classes. A total of 193,262 particles were imported and curated in RELION using 3D classification without alignment followed by Bayesian polishing with a box size of 300 pix (fourier cropped to 240 pix). Finally, the best-class consisting of 61,539 particles were imported and reconstructed in cryoSPARC using CTF refinement and non-uniform refinement, yielding a reconstruction with an overall resolution of 3.17 Å at 0.143 FSC criterion. In addition, the features of the reconstruction were improved following local refinement with a mask on the receptor resulting in a reconstruction with a nominal resolution of 3.71 Å. Since CXCL6 was not clearly discernible in the overall reconstruction, we prepared a composite map using the combine-focused-maps sub-module in Phenix (REF 28) with the overall reconstruction and the receptor-ligand focused map as inputs.

For the CXCL8-CXCR2-Go dataset, 2,152,291 particles were autopicked, extracted using a box size of 280 pix (fourier cropped to 70 pix), and subjected to heterogeneous refinement. Following re-extraction with a box size of 280 pix (fourier cropped to 180 pix), and subjected to heterogeneous refinement into three 3 classes. 99,138 particles corresponding to the best class following heterogeneous refinement was imported into RELION v4.0, subjected to Bayesian polishing with a box size of 300 pix (fourier cropped to 240 pix). The polished particles were then re-imported into cryoSPARC and was subjected to CTF refinement and non-uniform refinement to yield a map with a global resolution of 2.99 Å according to the gold-standard FSC cut-off of 0.143. Local refinement with a mask on the receptor and ligand was performed to yield a 3D reconstruction with a nominal resolution of 3.29 Å.

Local resolution of all maps were calculated using Blocres included within the cryoSPARC package^91^ with the half maps as input. Final maps were sharpened with phenix.auto_sharpen^92,93^ to enhance features for model building. Detailed pipelines for data processing and refinement are included in Supplementary Fig.

### Model building and refinement

The initial model of CXCR3 was generated from AlphaFold model (https://alphafold.ebi.ac.uk/entry/A0A0S2Z3W5), while the atomic coordinates of miniGo, and other component of G-protein (Gβ, Gγ and scFv16) were obtained from the cryo-EM structure of GALR1-miniGo complex^94^ (PDB: 7XJJ) and MT1-Gi complex^95^ (PDB: 7DB6), respectively. Ligand coordinates and geometric restraints were generated with Grade web server (Smart, O.S., Sharff A., Holstein, J., Womack, T.O., Flensburg, C., Keller, P., Paciorek, W., Vonrhein, C. and Bricogne G. (2021) Grade2 version 1.5.0. Cambridge, United Kingdom: Global Phasing Ltd.). These initial models were roughly docked into the density maps using UCSF ChimeraX^96,97^, followed by rigid body and flexible fitting of the coordinates with the jiggle fit and all atom refine module in COOT^98^. DeepEMhancer maps were used to fascilitate model building for low resolution region. The model so obtained was manually adjusted and rebuilt in COOT combined with iterative refinement with phenix.real_space_refine^93^ imposing secondary structural restraints. It is to be noted that although we prepared a complex of CXCR3 in presence of CXCL10, we could not observe any density for CXCL10, and therefore treated this structure as an apo state structure.

Coordinates of CXCR2 were generated in AlphaFold (https://alphafold.ebi.ac.uk/entry/P25025), while the atomic coordinates of miniGo, and other component of G-protein (Gβ, Gγ and scFv16) were obtained from the cryo-EM structure of EP54-C3aR-Go complex (PDB: 8I95). The initial model of the chemokines were obtained from the Swiss-model using previously solved CXCL8 structure as template (PDB: 6WZM). These initial models were docked into the individual EM maps with Chimera^96,97^, followed by flexible fitting of the docked models with the “all atom refine” module in COOT. The models so obtained were refined with phenix.real_space_refinement with secondary structural restraints against the EM maps after several rounds of manual readjustment in COOT. The final models were evaluated using Molprobity and the ‘‘Comprehensive Validation (cryo-EM)’’ sub-module within Phenix. Data collection, processing, and model refinement statistics are included in Extended Data Table 10. All figures in the manuscript were prepared using either Chimera or ChimeraX packages^96,97^.

## Data availability

All the data are included in the manuscript and any additional information required to reanalyse the data reported in this paper is available from the corresponding author upon reasonable request.

## Code availability

The cryo-EM structures are deposited in Protein Data Bank (PDB) and Electron Microscopy Data Bank (EMDB) with accession numbers 8XWA and EMD-38732 for CXCL1-CXCR2-Go (Receptor-Ligand Focused), 8XWV and EMD-38743 for CXCL1-CXCR2-Go (Overall), 8XVU and EMD-38719 for CXCL2-CXCR2-Go (Receptor-Ligand Focused), 8XXH and EMD-38749 for CXCL2-CXCR2-Go (Overall), 8XWF and EMD-38734 for CXCL3-CXCR2-Go (Receptor-Ligand Focused), 8XX3 and EMD-38744 for CXCL3-CXCR2-Go (Overall), 8XWS and EMD-38742 for CXCL5-CXCR2-Go (Receptor-Ligand Focused), 8XX7 and EMD-38748 for CXCL5-CXCR2-Go (Overall), 8XWM and EMD-38738 for CXCL6-CXCR2-Go (Receptor-Ligand Focused), 8XXR and EMD-38759 for CXCL6-CXCR2-Go (Overall), 8XXX and EMD-38764 for CXCL6-CXCR2-Go (composite), 8XWN and EMD-38739 for CXCL8-CXCR2-Go (Receptor-Ligand Focused), 8XX6 and EMD-38747 for CXCL8-CXCR2-Go (Overall), 8XXY and EMD-38765 for Apo-CXCR3-Go (Receptor-Ligand Focused), 8XXZ and EMD-38766 for Apo-CXCR3-Go (Overall), 8Y0H and EMD-38803 for VUF11418-CXCR3-Go (Receptor-Ligand Focused), 8Y0N and EMD-38809 for VUF11418-CXCR3-Go (Overall), 8XYI and EMD-38774 for VUF10661-CXCR3-Go (Receptor-Ligand Focused), 8XYK and EMD-38776 for VUF10661-CXCR3-Go (Overall). Source data are provided with this paper. This paper does not report any original code.

## Acknowledgement

Research in A.K.S.’s laboratory is supported by the Senior Fellowship of the DBT Wellcome Trust India Alliance (IA/S/20/1/504916) awarded to A.K.S., the Science and Engineering Research Board (SPR/2020/000408 and IPA/2020/000405), the Indian Council of Medical research (F.NO.52/15/2020/BIO/BMS), and IIT Kanpur. A.K.S. is Sonu Agrawal Memorial Chair Professor. S.S. is funded by the Prime Minister’s Research Fellowship (PMRF). This work was supported by grants from the JSPS KAKENHI, grant numbers 21H05037 (O.N.), 22K19371 and 22H02751 (W.S.), and 23KJ0491 (F.K.S.); the Kao Foundation for Arts and Sciences (W.S.); the Takeda Science Foundation (W.S.); the Lotte Foundation (W.S.); and the Platform Project for Supporting Drug Discovery and Life Science Research [Basis for Supporting Innovative Drug Discovery and Life Science Research (BINDS)] from the Japan Agency for Medical Research and Development (AMED), grant numbers JP22ama121012 (O.N.) and JP22ama121002 (support number 3272; O.N.). AC was supported by the Luxembourg Institute of Health (LIH) through the NanoLux Platform, the Luxembourg National Research Fund (INTER/FNRS grants INTER 20/15084569 and CORE C23/BM/18068832) and the F.R.S.-FNRS-Télévie (grants 7.8504.20, 7.4502.21 and 7.8508.22).

## Authors’ contribution

SS and SSh reconstituted the complexes for structural analysis with help from SSa, SM, AD, SMo and MKY, and carried out the functional assays on CXCR2 and CXCR3 together with NZ; FKS prepared and screened the cryo-EM grids, collected and processed the cryo-EM data, and solved the structures with help from HA, TK, and YI; MG and RB refined and analyzed the structures and prepared the figures; RL provided small molecule agonists of CXCR3; AC carried out and analyzed the chemokine profiling experiments; RB, WS, ON and AKS supervised the overall study.

