## Supplemental Material for "Molecular basis of ligand promiscuity, structural mimicry, and atypical dimerization in the chemokine receptors"

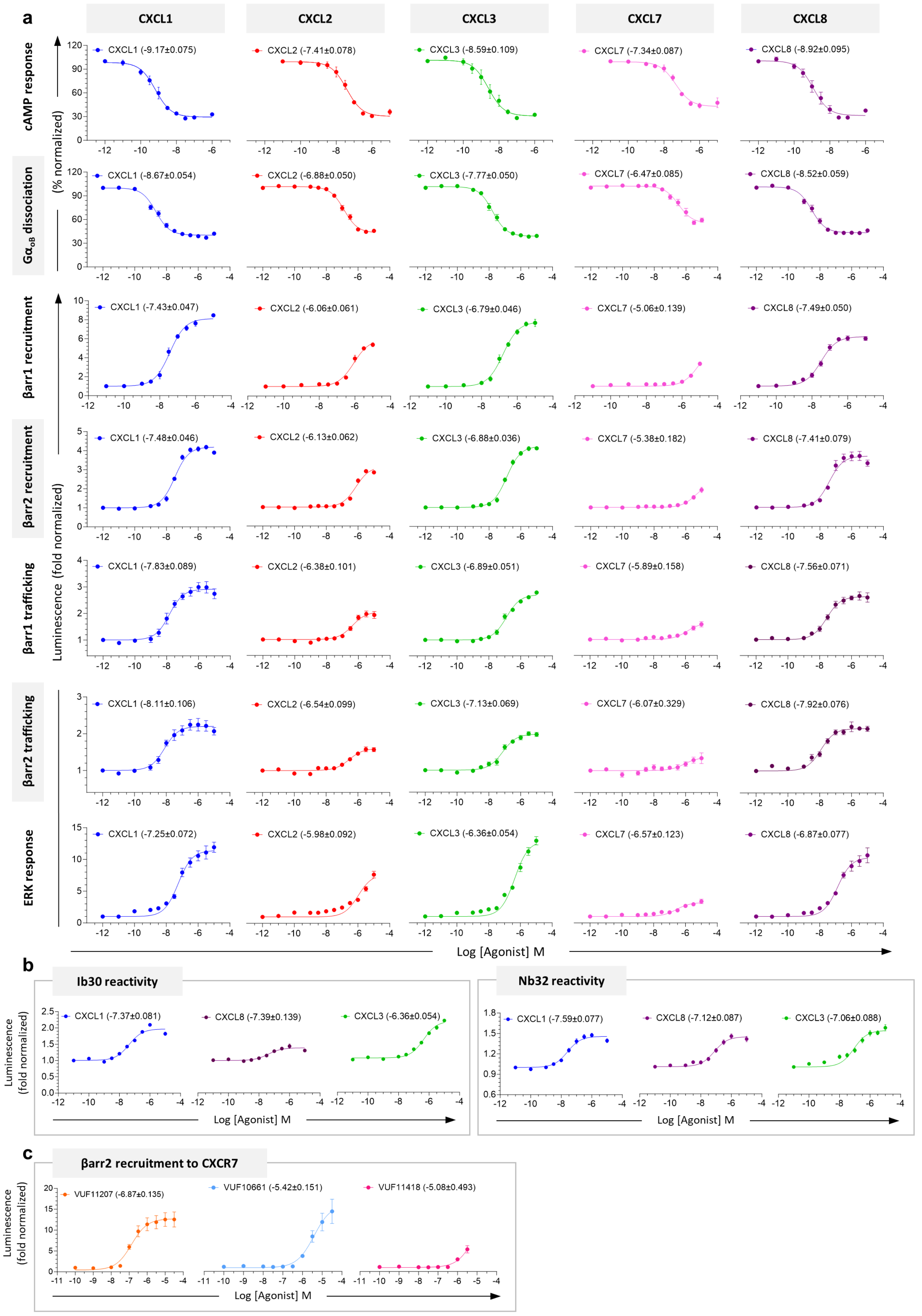
a
CXCL1
CXCL2
CXCL3
CXCL7
CXCL8
cAMP response
GαoB dissociation
(% normalized)
CXCL1 (-9.17±0.075)
CXCL2 (-7.41±0.078)
CXCL3 (-8.59±0.109)
CXCL7 (-7.34±0.087)
CXCL8 (-8.92±0.095)
CXCL1 (-8.67±0.054)
CXCL2 (-6.88±0.050)
CXCL3 (-7.77±0.050)
CXCL7 (-6.47±0.085)
CXCL8 (-8.52±0.059)
βarr1 recruitment
βarr2 recruitment
βarr1 trafficking
βarr2 trafficking
ERK response
Luminescence (fold normalized)
CXCL1 (-7.43±0.047)
CXCL2 (-6.06±0.061)
CXCL3 (-6.79±0.046)
CXCL7 (-5.06±0.139)
CXCL8 (-7.49±0.050)
CXCL1 (-7.48±0.046)
CXCL2 (-6.13±0.062)
CXCL3 (-6.88±0.036)
CXCL7 (-5.38±0.182)
CXCL8 (-7.41±0.079)
CXCL1 (-7.83±0.089)
CXCL2 (-6.38±0.101)
CXCL3 (-6.89±0.051)
CXCL7 (-5.89±0.158)
CXCL8 (-7.56±0.071)
CXCL1 (-8.11±0.106)
CXCL2 (-6.54±0.099)
CXCL3 (-7.13±0.069)
CXCL7 (-6.07±0.329)
CXCL8 (-7.92±0.076)
CXCL1 (-7.25±0.072)
CXCL2 (-5.98±0.092)
CXCL3 (-6.36±0.054)
CXCL7 (-6.57±0.123)
CXCL8 (-6.87±0.077)
Log [Agonist] M
b
Ib30 reactivity
Nb32 reactivity
Luminescence (fold normalized)
CXCL1 (-7.37±0.081)
CXCL8 (-7.39±0.139)
CXCL3 (-6.36±0.054)
CXCL1 (-7.59±0.077)
CXCL8 (-7.12±0.087)
CXCL3 (-7.06±0.088)
Log [Agonist] M
c
βarr2 recruitment to CXCR7
Luminescence (fold normalized)
VUF11207 (-6.87±0.135)
VUF10661 (-5.42±0.151)
VUF11418 (-5.08±0.493)
Log [Agonist] M

**Extended Data Fig. 1:**

**a-b,** Dose curves of response elicited by CXCR2 downstream to stimulation with different agonists were measured using a multitude of assays. Data (mean±SEM) represents three-six independent biological replicates, performed in duplicate, and normalized with respect to signal observed at lowest dose, treated either as 100% (for cAMP response and GoB dissociation), or 1 (βarr1/2 recruitment, βarr1/2 trafficking, ERK assay, Ib30 reactivity and Nb32 reactivity). **c,** βarr2 recruitment to CXCR7 as measured by TANGO assay confirms the dual agonistic property of VUF10661 and VUF11418. Data (mean±SEM) represents three independent biological replicates, performed in duplicate, and normalized with respect to signal observed at lowest dose, treated as 1.

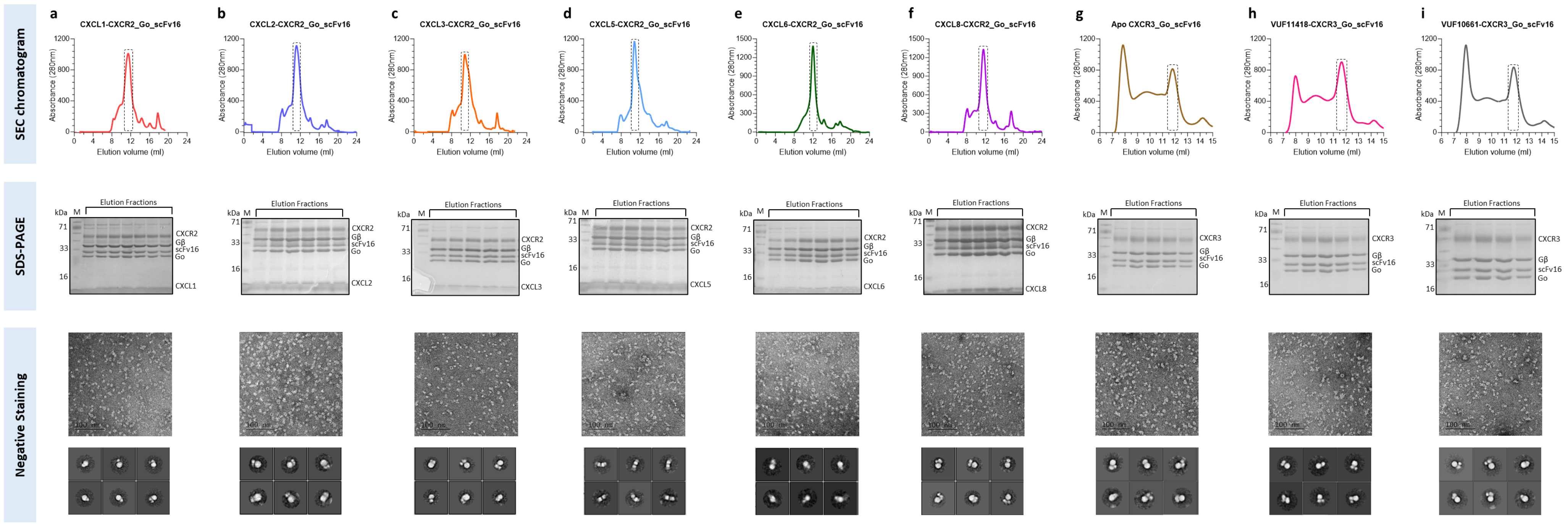


**Extended Data Fig. 2: Purification of CXCR2 and CXCR3 complexes and visualization through negative-staining EM.**

**a-i,** Size exclusion chromatography profile, SDS-PAGE and negative staining-EM of CXCL1-CXCR2-Go-scFv16, CXCL2-CXCR2-Go-scFv16, CXCL3-CXCR2-Go-scFv16, CXCL5-CXCR2-Go-scFv16, CXCL6-CXCR2-Go-scFv16, CXCL8-CXCR2-Go-scFv16, Apo-CXCR3-Go-scFv16, VUF11418-CXCR3-Go-scFv16 and VUF10661-CXCR3-Go-scFv16 complexes, respectively.

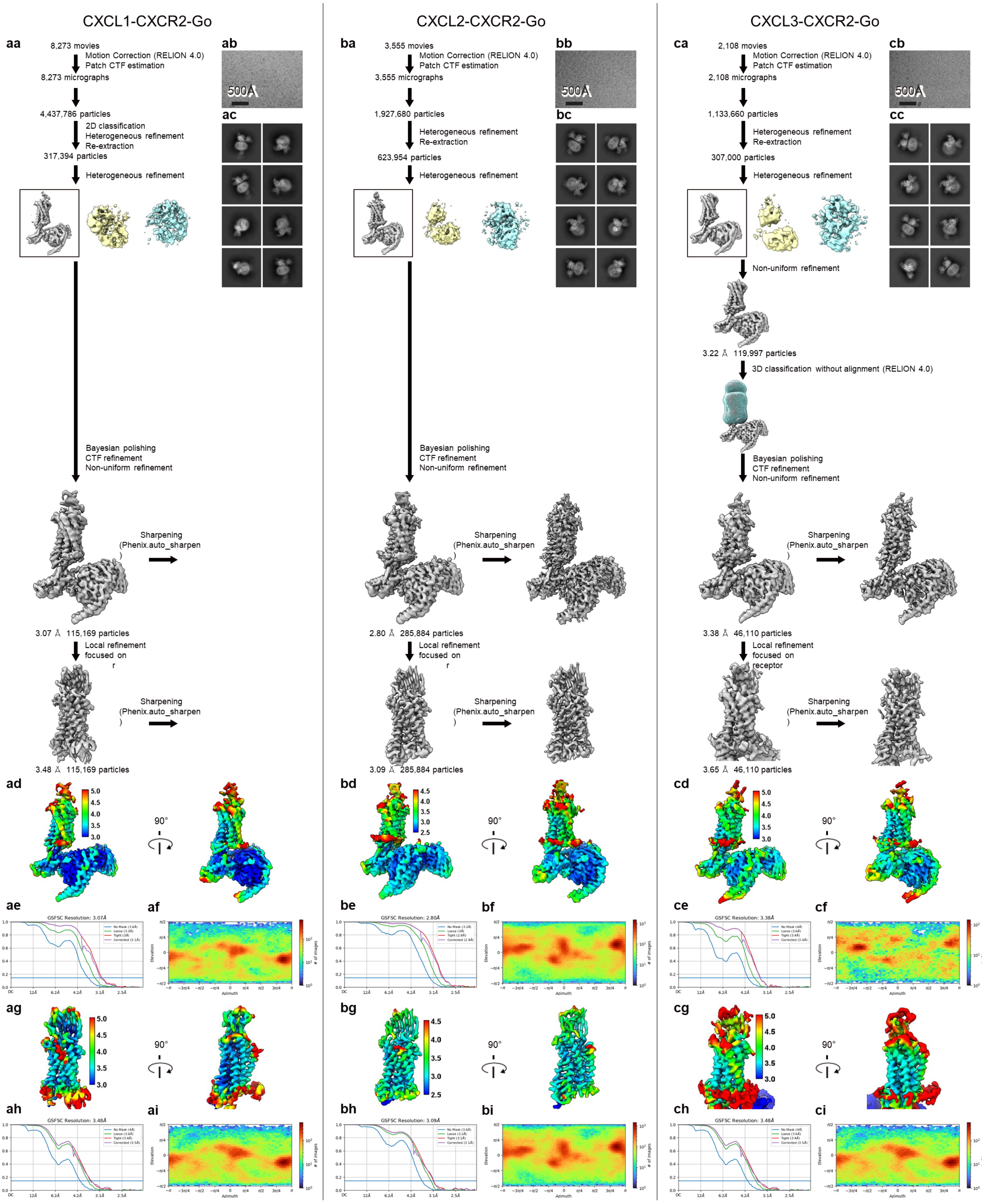

CXCL1-CXCR2-Go
CXCL2-CXCR2-Go
CXCL3-CXCR2-Go
aa
8,273 movies
Motion Correction (RELION 4.0)
Patch CTF estimation
8,273 micrographs
4,437,786 particles
2D classification
Heterogeneous refinement
Re-extraction
317,394 particles
Heterogeneous refinement
Bayesian polishing
CTF refinement
Non-uniform refinement
Sharpening
(Phenix.auto_sharpen
)
3.07 Å 115,169 particles
Local refinement
focused on
r
Sharpening
(Phenix.auto_sharpen
)
3.48 Å 115,169 particles
ab
500Å
ac
ba
3,555 movies
Motion Correction (RELION 4.0)
Patch CTF estimation
3,555 micrographs
1,927,680 particles
Heterogeneous refinement
Re-extraction
623,954 particles
Heterogeneous refinement
Bayesian polishing
CTF refinement
Non-uniform refinement
Sharpening
(Phenix.auto_sharpen
)
2.80 Å 285,884 particles
Local refinement
focused on
r
Sharpening
(Phenix.auto_sharpen
)
3.09 Å 285,884 particles
bb
500Å
bc
ca
2,108 movies
Motion Correction (RELION 4.0)
Patch CTF estimation
2,108 micrographs
1,133,660 particles
Heterogeneous refinement
Re-extraction
307,000 particles
Heterogeneous refinement
Non-uniform refinement
3.22 Å 119,997 particles
3D classification without alignment (RELION 4.0)
Bayesian polishing
CTF refinement
Non-uniform refinement
Sharpening
(Phenix.auto_sharpen
)
3.38 Å 46,110 particles
Local refinement
focused on
receptor
Sharpening
(Phenix.auto_sharpen
)
3.65 Å 46,110 particles
cb
500Å
cc
ad
5.0
4.5
4.0
3.5
3.0
90°
bd
4.5
4.0
3.5
3.0
2.5
90°
cd
5.0
4.5
4.0
3.5
3.0
90°
ae
GSFSC Resolution: 3.07Å
af
be
GSFSC Resolution: 2.80Å
bf
ce
GSFSC Resolution: 3.38Å
cf
ag
5.0
4.5
4.0
3.5
3.0
90°
bg
4.5
4.0
3.5
3.0
2.5
90°
cg
5.0
4.5
4.0
3.5
3.0
90°
ah
GSFSC Resolution: 3.48Å
ai
bh
GSFSC Resolution: 3.09Å
bi
ch
GSFSC Resolution: 3.48Å
ci

**Extended Data Fig. 3: Cryo-EM data processing pipeline of CXCL1-CXCR2-Go, CXCL2-CXCR2-Go and CXCL3-CXCR2-Go.**

**aa, ba, ca,** Schematic representations of the cryo-EM data processing workflow for CXCL1-CXCR2-Go (**aa**), CXCL2-CXCR2-Go (**ba**) and CXCL3-CXCR2-Go (**ca**).

**ab, bb, cb,** Representative cryo-EM images of the CXCL1-CXCR2-Go (**ab**), CXCL2-CXCR2-Go (**bb**) and CXCL3-CXCR2-Go (**cb**) recorded on a 300 kV Titan Krios with a K3 camera.

**ac, bc, cc,** Representative 2D averages of curated particles for CXCL1-CXCR2-Go (**ac**), CXCL2-CXCR2-Go (**bc**) and CXCL3-CXCR2-Go (**cc**).

**ad, bd, cd,** Local resolution of the overall refined maps for CXCL1-CXCR2-Go (**ad**), CXCL2-CXCR2-Go (**bd**) and CXCL3-CXCR2-Go (**cd**).

**ae, be, ce,** Gold standard fourier shell correlation curve (FSC) at 0.143 threshold for overall refined maps of CXCL1-CXCR2-Go (**ae**), CXCL2-CXCR2-Go (**be**) and CXCL3-CXCR2-Go (**ce**).

**af, bf, cf,** Angular distribution of the overall refinement for CXCL1-CXCR2-Go (**af**), CXCL2-CXCR2-Go (**bf**) and CXCL3-CXCR2-Go (**cf**).

**ag, bg, cg,** Local resolution of the local refined maps for CXCL1-CXCR2-Go (**ag**), CXCL2-CXCR2-Go (**bg**) and CXCL3-CXCR2-Go (**cg**).

**ah, bh, ch,** Gold standard fourier shell correlation curve (FSC) at 0.143 threshold for local refined maps of CXCL1-CXCR2-Go (**ah**), CXCL2-CXCR2-Go (**bh**) and CXCL3-CXCR2-Go (**ch**).

**ai, bi, ci,** Angular distribution of the local refinement for CXCL1-CXCR2-Go (**ai**), CXCL2-CXCR2-Go (**bi**) and CXCL3-CXCR2-Go (**ci**).

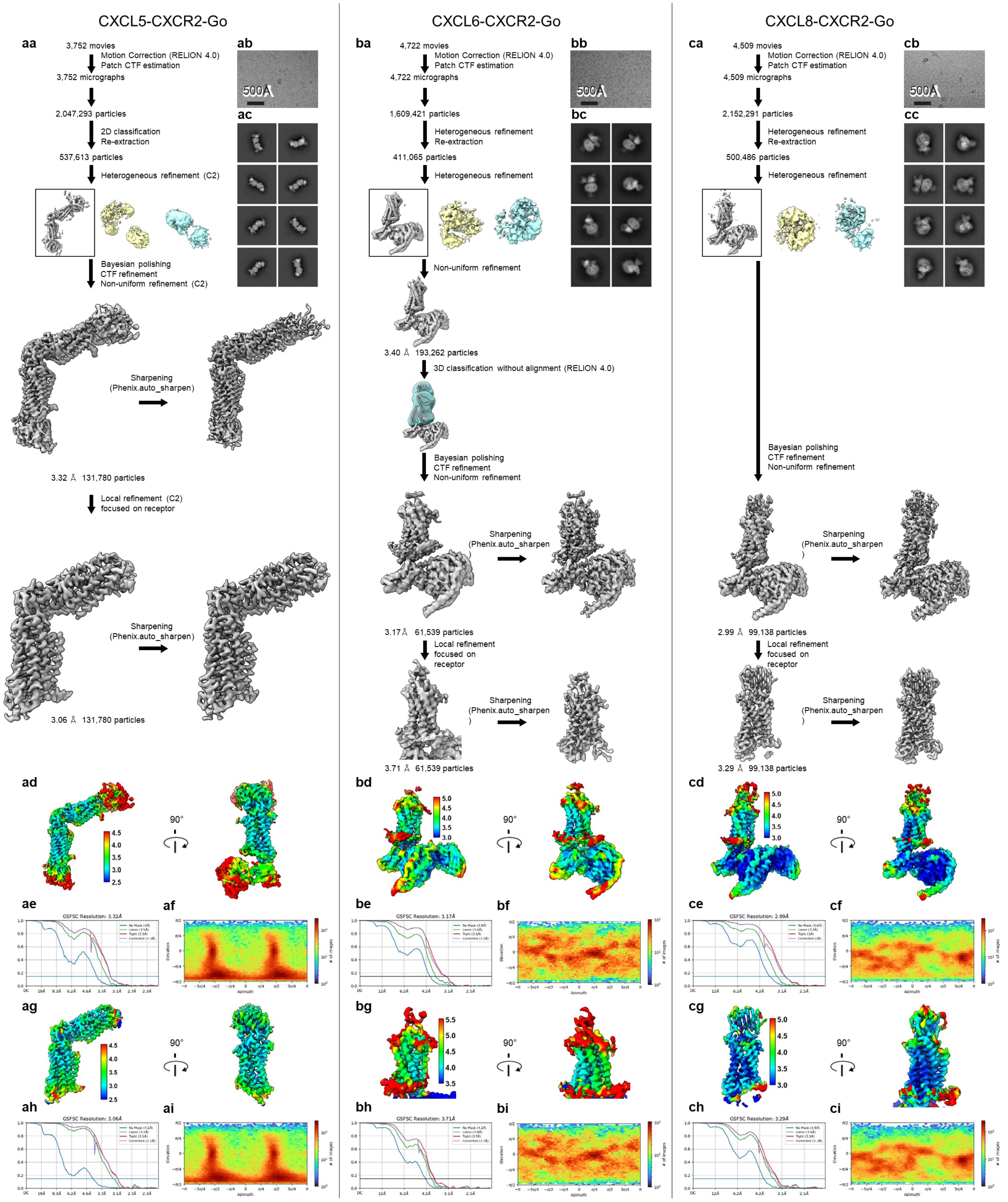

CXCL5-CXCR2-Go
CXCL6-CXCR2-Go
CXCL8-CXCR2-Go
aa
3,752 movies
Motion Correction (RELION 4.0)
Patch CTF estimation
3,752 micrographs
2,047,293 particles
2D classification
Re-extraction
537,613 particles
Heterogeneous refinement (C2)
Bayesian polishing
CTF refinement
Non-uniform refinement (C2)
Sharpening
(Phenix.auto_sharpen)
3.32 Å 131,780 particles
Local refinement (C2)
focused on receptor
3.06 Å 131,780 particles
ab
500Å
ac
ba
4,722 movies
Motion Correction (RELION 4.0)
Patch CTF estimation
4,722 micrographs
1,609,421 particles
Heterogeneous refinement
Re-extraction
411,065 particles
Heterogeneous refinement
Non-uniform refinement
3.40 Å 193,262 particles
3D classification without alignment (RELION 4.0)
Bayesian polishing
CTF refinement
Non-uniform refinement
3.17 Å 61,539 particles
Local refinement
focused on
receptor
3.71 Å 61,539 particles
bb
bc
ca
4,509 movies
Motion Correction (RELION 4.0)
Patch CTF estimation
4,509 micrographs
2,152,291 particles
Heterogeneous refinement
Re-extraction
500,486 particles
Heterogeneous refinement
Bayesian polishing
CTF refinement
Non-uniform refinement
2.99 Å 99,138 particles
Local refinement
focused on
receptor
3.29 Å 99,138 particles
cb
cc
ad
90°
bd
cd
ae
GSFSC Resolution: 3.32Å
af
be
GSFSC Resolution: 3.17Å
bf
ce
GSFSC Resolution: 2.99Å
cf
ag
bg
cg
ah
GSFSC Resolution: 3.06Å
ai
bh
GSFSC Resolution: 3.71Å
bi
ch
GSFSC Resolution: 3.29Å
ci
Azimuth
Elevation

**Extended Data Fig. 4: Cryo-EM data processing pipeline of CXCL5-CXCR2-Go, CXCL6-CXCR2-Go and CXCL8-CXCR2-Go.**

**aa, ba, ca,** Schematic representations of the cryo-EM data processing workflow for CXCL5-CXCR2-Go (**aa**), CXCL6-CXCR2-Go (**ba**) and CXCL8-CXCR2-Go (**ca**).

**ab, bb, cb,** Representative cryo-EM images of the CXCL5-CXCR2-Go (**ab**), CXCL6-CXCR2-Go (**bb**) and CXCL8-CXCR2-Go (**cb**) recorded on a 300 kV Titan Krios with a K3 camera.

**ac, bc, cc,** Representative 2D averages of curated particles for CXCL5-CXCR2-Go (**ac**), CXCL6-CXCR2-Go (**bc**) and CXCL8-CXCR2-Go (**cc**).

**ad, bd, cd,** Local resolution of the overall refined maps for CXCL5-CXCR2-Go (**ad**), CXCL6-CXCR2-Go (**bd**) and CXCL8-CXCR2-Go (**cd**).

**ae, be, ce,** Gold standard fourier shell correlation curve (FSC) at 0.143 threshold for overall refined maps of CXCL5-CXCR2-Go (**ae**), CXCL6-CXCR2-Go (**be**) and CXCL8-CXCR2-Go (**ce**).

**af, bf, cf,** Angular distribution of the overall refinement for CXCL5-CXCR2-Go (**af**), CXCL6-CXCR2-Go (**bf**) and CXCL8-CXCR2-Go (**cf**).

**ag, bg, cg,** Local resolution of the local refined maps for CXCL5-CXCR2-Go (**ag**), CXCL6-CXCR2-Go (**bg**) and CXCL8-CXCR2-Go (**cg**).

**ah, bh, ch,** Gold standard fourier shell correlation curve (FSC) at 0.143 threshold for local refined maps of CXCL5-CXCR2-Go (**ah**), CXCL6-CXCR2-Go (**bh**) and CXCL8-CXCR2-Go (**ch**).

**ai, bi, ci,** Angular distribution of the local refinement for CXCL5-CXCR2-Go (**ai**), CXCL6-CXCR2-Go (**bi**) and CXCL8-CXCR2-Go (**ci**).

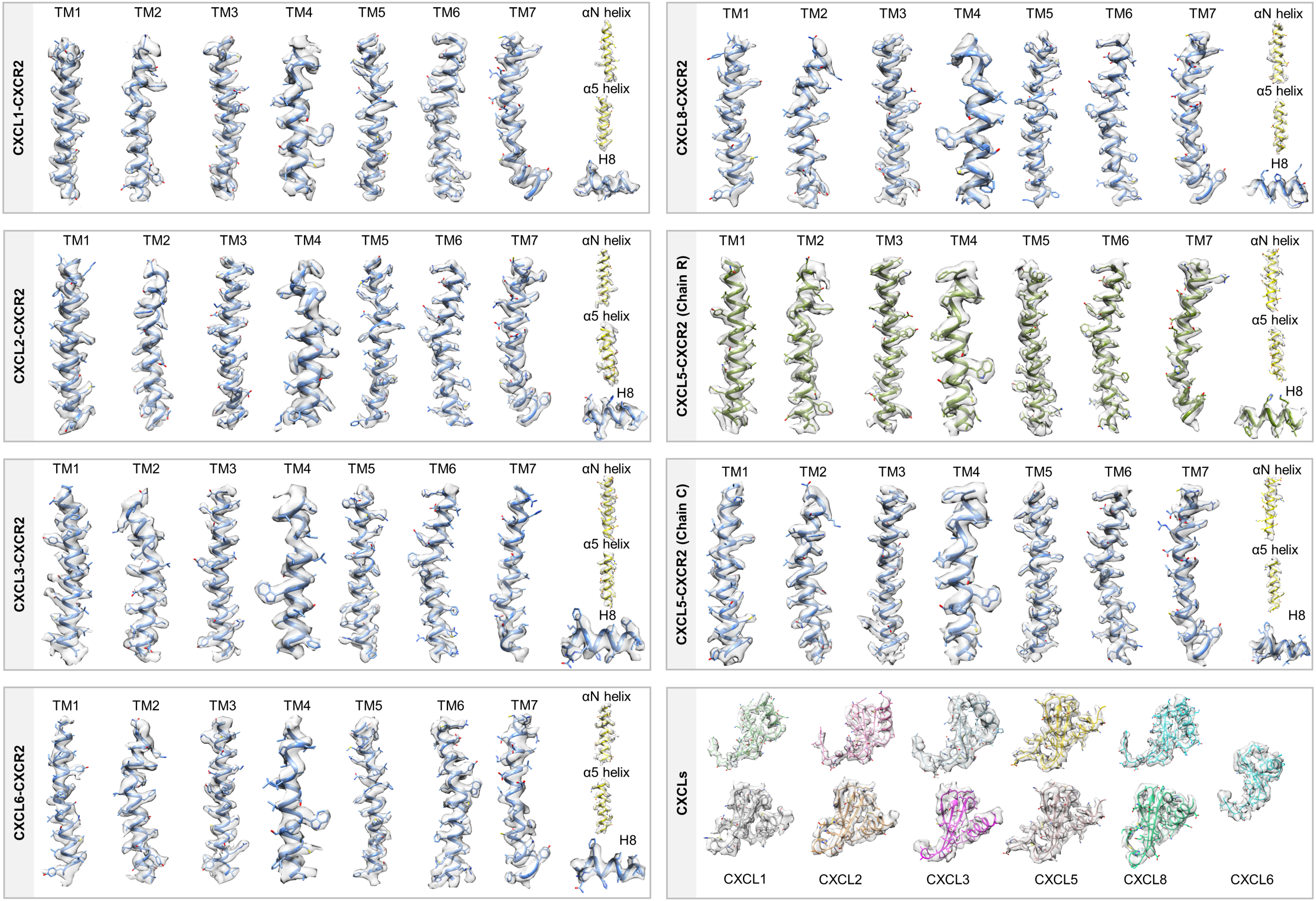

CXCL1-CXCR2
TM1
TM2
TM3
TM4
TM5
TM6
TM7
αN helix
α5 helix
H8
CXCL8-CXCR2
CXCL2-CXCR2
CXCL5-CXCR2 (Chain R)
CXCL3-CXCR2
CXCL5-CXCR2 (Chain C)
CXCL6-CXCR2
CXCLs
CXCL1
CXCL2
CXCL3
CXCL5
CXCL8
CXCL6

**Extended Data Fig. 5: Exemplary electron density maps of the CXCR2 complexes.**

EM densities for the TMs, helix 8, αN helix and α5 helix of CXCR2 structures and CXCL1, CXCL2, CXCL3, CXCL5, CXCL6 and CXCL8.

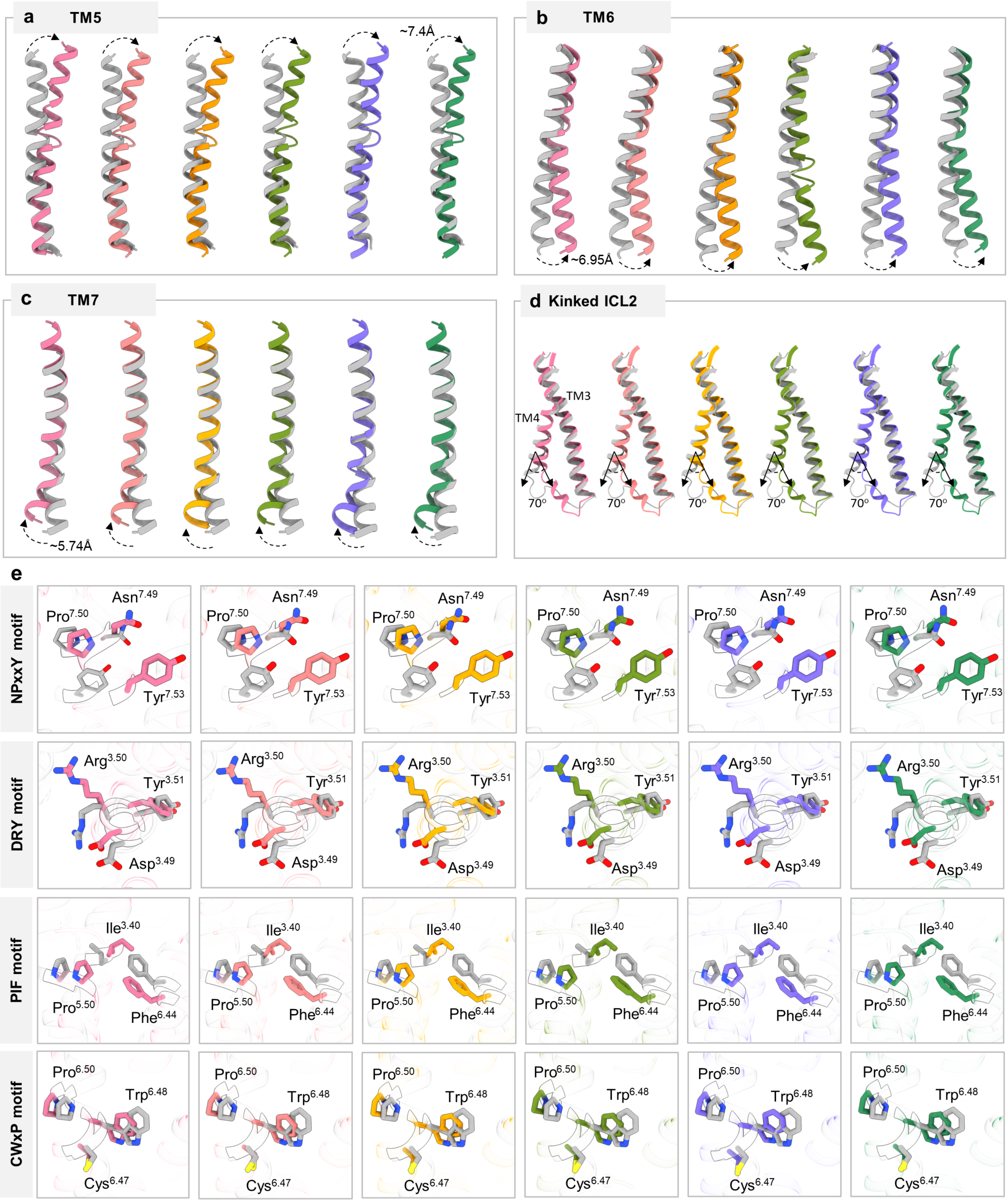

a TM5
~7.4Å
b TM6
~6.95Å
c TM7
~5.74Å
d Kinked ICL2
TM3
TM4
70°
e
NPxxY motif
Asn7.49
Pro7.50
Tyr7.53
DRY motif
Arg3.50
Tyr3.51
Asp3.49
PIF motif
Ile3.40
Pro5.50
Phe6.44
CWxP motif
Pro6.50
Trp6.48
Cys6.47

**Extended Data Fig. 6: TM movements, kink formation in ICL2 and conformational changes in conserved motifs.**

**a-c,** Activation dependent changes in TMs of chemokine bound CXCR2 structures compared with the inactive state structure (PDB:6LFL, gray). The dotted arrows indicate direction of movement from the inactive to active state. The respective degrees of movements in the corresponding TMs have been mentioned. **d,** Formation of a kink in ICL2 at the junction of TM3 and TM4 upon activation of CXCR2. **e,** Conformational changes in the conserved microswitches of CXCR2: NPxxY, DRY, PIF and CWxP (pink: CXCL1 bound CXCR2, salmon: CXCL2 bound CXCR2, yellow: CXCL3 bound CXCR2, olive green: CXCL5 bound CXCR2, purple: CXCL6 bound CXCR2, dark green: CXCL8 bound CXCR2).

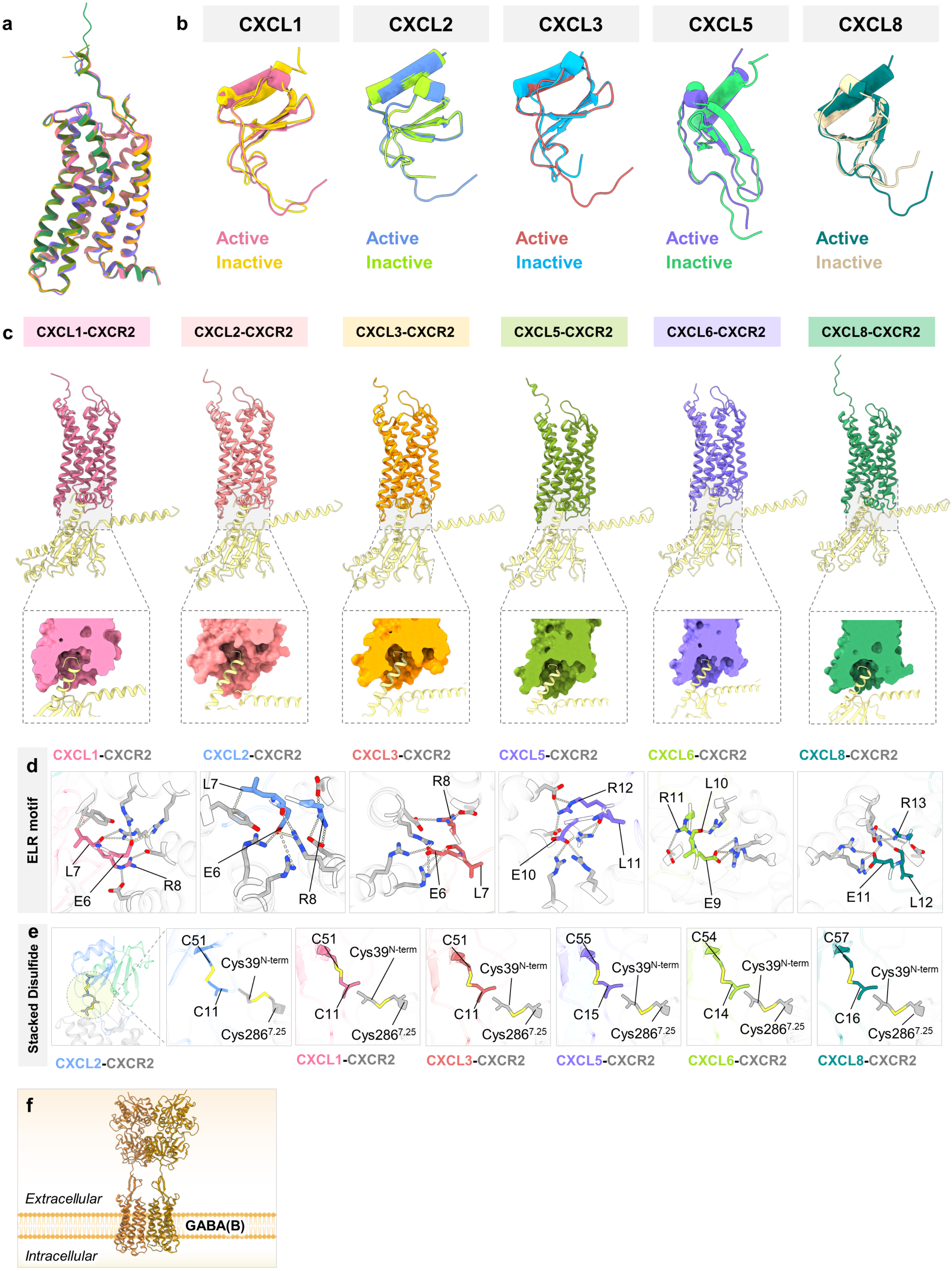
a
b
CXCL1
CXCL2
CXCL3
CXCL5
CXCL8
Active
Inactive
c
CXCL1-CXCR2
CXCL2-CXCR2
CXCL3-CXCR2
CXCL5-CXCR2
CXCL6-CXCR2
CXCL8-CXCR2
d
ELR motif
L7
E6
R8
R12
E10
L11
R11
L10
E9
R13
E11
L12
e
Stacked Disulfide
C51
C11
C55
C15
C54
C14
C57
C16
Cys39N-term
Cys2867.25
f
Extracellular
GABA(B)
Intracellular

**Extended Data Fig. 7: Structural superimposition and critical interactions at the receptor-G protein and receptor-ligand interface. a,** Overall RMSD among the different CXCR2 structures upon superimposition is less than 0.45Å. **b,** Structural alignment depicting the formation of an extended "U-shaped" loop at the N-terminus of the chemokines upon binding to CXCR2. **c,** α5 helix of Gαo docks into the cytoplasmic cavity of CXCR2. Only receptor and Gαo have been shown to directly visualize the interaction of Gαo with CXCR2. **(In inset)** Surface slice representation of the receptor along with ribbon diagram of Gαo. **d-e,** Magnified view of the interactions in ELR motif **(d)** and stacked disulfide between chemokine and receptor **(e)** for the different CXCR2 complexes. Ionic interactions have been depicted as gray dashed lines. **f,** Structural representation of dimeric GABA(B) receptor(PDB: 7C7Q), a prototypical class C GPCR.

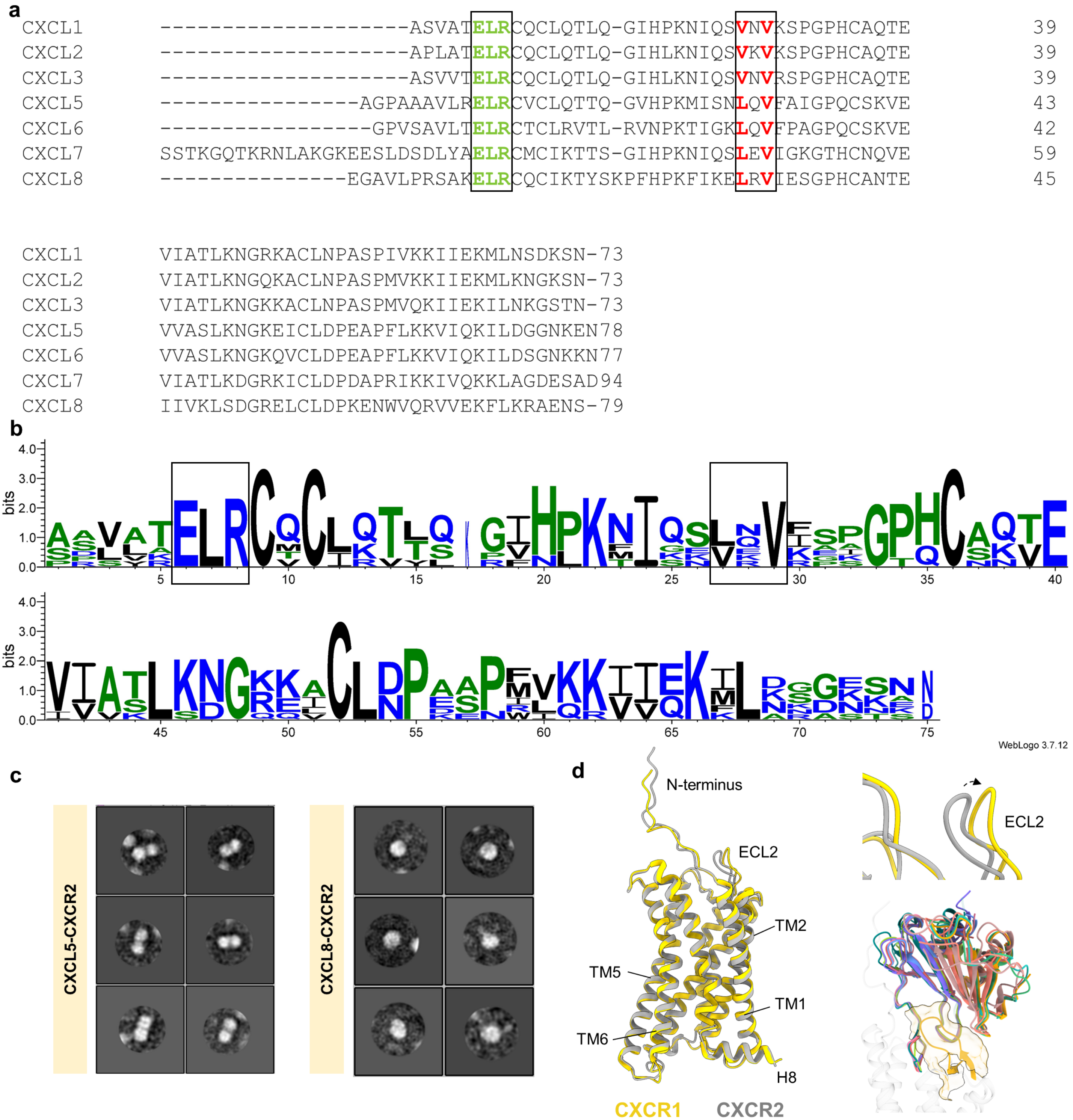


**Extended Data Fig. 8: Sequence alignment of all CXCR2 binding CXCLs, negative staining showing CXCL5 induced dimerization and ECL2 movement in CXCR1.**

**a-b,** Sequence alignment of all CXCR2 binding CXCLs, highlighting the conserved ELR motif and hydrophobic patch that mediates ligand dimerization. Sequence alignment was performed using Clustal Omega (https://www.ebi.ac.uk/jdispatcher/msa/clustalo) and subsequently the consensus sequence logo was generated using the WEBLOGO tool (https://weblogo.threeplusone.com/create.cgi). **c,** Negative staining 2D class averages of CXCL5-CXCR2 and CXCL8-CXCR2 complexes**. d,** Superimposition of CXCL8 bound CXCR1 (PDB: 8IC0, yellow) and CXCL8 bound CXCR2 (grey) highlights the outward movement of ECL2 in CXCR1 (Left and Right top). ECL2 of CXCR1 exhibits steric clash with the second protomer of CXCLs (Right bottom).

```
CXCL1     --------------------ASVATELRCQCLQTLQG-IHP--KNIQSVNV-KSPGPHCA     36
CXCL2     --------------------APLATELRCQCLQTLQG-IHL--KNIQSVKV-KSPGPHCA     36
CXCL3     --------------------ASVVTELRCQCLQTLQG-IHL--KNIQSVNV-RSPGPHCA     36
CXCL4     -------------------EAEEDGDLQCLCVKTTSQ-VRP--RHITSLEV-IKAGPHCP     37
CXCL5     ----------------AGPAAAVLRELRCVCLQTTQG-VHP--KMISNLQV-FAIGPQCS     40
CXCL6     -----------------GPVSAVLTELRCTCLRVTLR-VNP--KTIGKLQV-FPAGPQCS     39
CXCL7     SSTKGQTKRNLAKGKEESLDSDLYAELRCMCIKTTSG-IHP--KNIQSLEV-IGKGTHCN     56
CXCL8     ---------------EGAVLPRSAKELRCQCIKTYSKPFHP--KFIKELRV-IESGPHCA     42
CXCL9     --------------------TPVVRKGRCSCISTNQGTIHL--QSLKDLKQ-FAPSPSCE     37
CXCL10    --------------------VPLSRTVRCTCISISNQPVNP--RSLEKLEI-IPASQFCP     37
CXCL11    --------------------FPMFKRGRCLCIGPGVKAVKV--ADIEKASI-MYPSNNCD     37
CXCL12    --------------------KPVSLSYRCPCRFFESH-VAR--ANVKHLKI-LN-TPNCA     35
CXCL13    ------------------VLEVYYTSLRCRCVQESSVFIPR--RFIDRIQI-LPRGNGCP     39
CXCL14    --------------------------SKCKCSRKGPK-IRY--SDVKKLEM-KPKYPHCE     30
CXCL16    --------------------NEGSVTGSCYCGKRISSDSPPSVQFMNRLRKHLRAYHRCL     40

CXCL1     ---QTEVIATLK-----NGRKACLNPASPIVKKIIEKMLNSDKSN---------------     73
CXCL2     ---QTEVIATLK-----NGQKACLNPASPMVKKIIEKMLKNGKSN---------------     73
CXCL3     ---QTEVIATLK-----NGKKACLNPASPMVQKIIEKILNKGSTN---------------     73
CXCL4     ---TAQLIATLK-----NGRKICLDLQAPLYKKIIKKLLES-------------------     70
CXCL5     ---KVEVVASLK-----NGKEICLDPEAPFLKKVIQKILDGGNKEN--------------     78
CXCL6     ---KVEVVASLK-----NGKQVCLDPEAPFLKKVIQKILDSGNKKN--------------     77
CXCL7     ---QVEVIATLK-----DGRKICLDPDAPRIKKIVQKKLAGDESAD--------------     94
CXCL8     ---NTEIIVKLS-----DGRELCLDPKENWVQRVVEKFLKRAENS---------------     79
CXCL9     ---KIEIIATLK-----NGVQTCLNPDSADVKELIKKWEKQVSQKKKQKNGKKHQKKKVL     89
CXCL10    ---RVEIIATMKK----KGEKRCLNPESKAIKNLLKAVSKERSKRSP-------------     77
CXCL11    ---KIEVIITLKE----NKGQRCLNPKSKQARLIIKKVERKNF-----------------     73
CXCL12    ----LQIVARLKN----NNRQVCIDPKLKWIQEYLEKALNK-------------------     68
CXCL13    ---RKEIIVWKK-----NKSIVCVDPQAEWIQRMMEVLRKRSSSTLPVPVFKRKIP----     87
CXCL14    ---EKMVIITTKSVSRYRGQEHCLHPKLQSTKRFIKWYNAWNEKRRVYEE----------     77
CXCL16    YYTRFQLL----------SWSVCGGNKDPWVQELMSCLDLKECG-HAYSGIVAHQKHLLP     89

CXCL1     --------------     73
CXCL2     --------------     73
CXCL3     --------------     73
CXCL4     --------------     70
CXCL5     --------------     78
CXCL6     --------------     77
CXCL7     --------------     94
CXCL8     --------------     79
CXCL9     KVRKSQRSRQKKTT     103
CXCL10    --------------     77
CXCL11    --------------     73
CXCL12    --------------     68
CXCL13    --------------     87
CXCL14    --------------     77
CXCL16    --------------     89
```

**Extended Data Fig. 9: Sequence alignment of all C-X-C type chemokine ligands.**

Sequence alignment of all C-X-C type chemokine ligands, highlighting the conserved ELR motif. Sequence alignment was performed using Clustal Omega (https://www.ebi.ac.uk/jdispatcher/msa/clustalo) and subsequently the consensus sequence logo was generated using the WEBLOGO tool (https://weblogo.threeplusone.com/create.cgi).

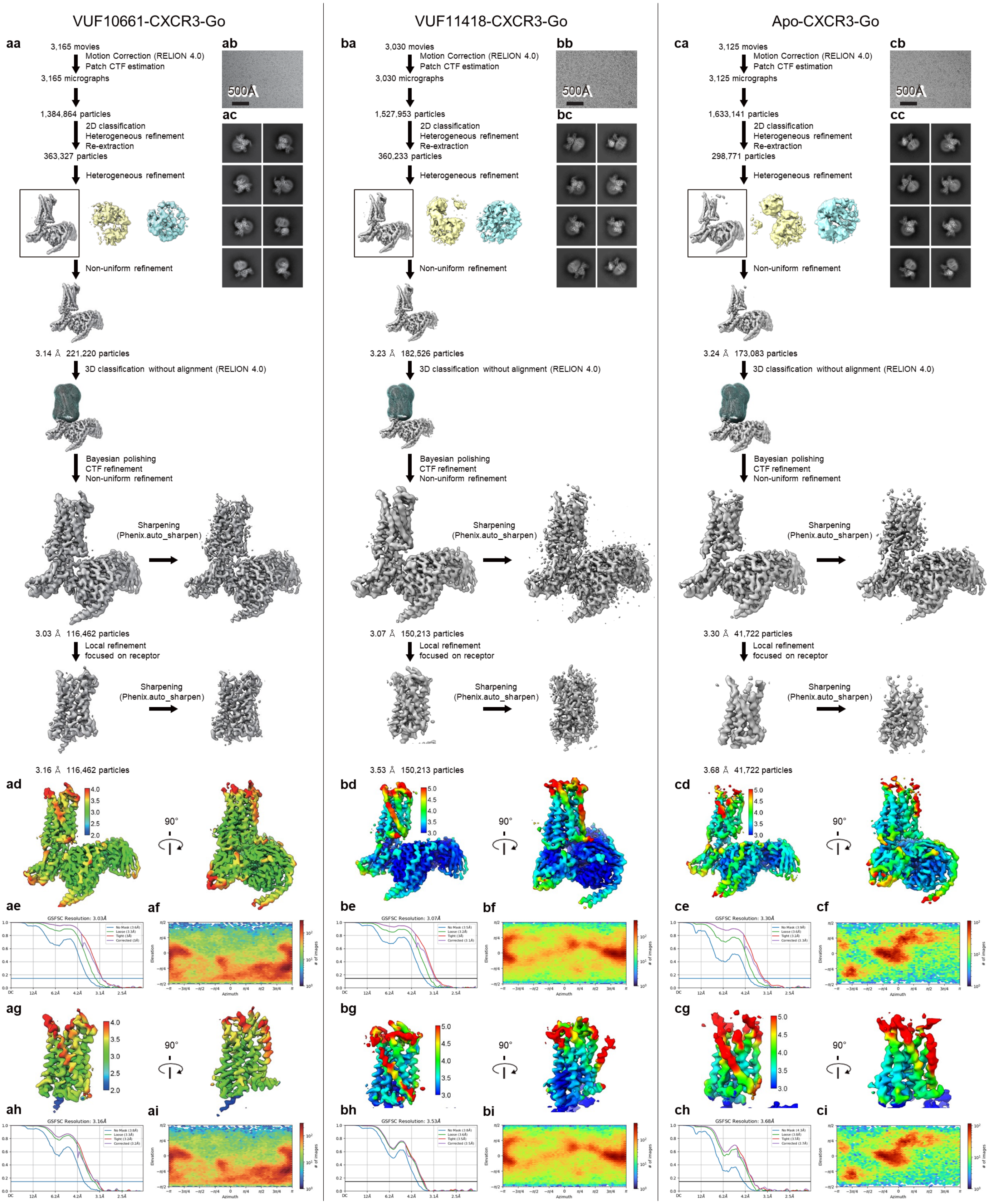

VUF10661-CXCR3-Go
VUF11418-CXCR3-Go
Apo-CXCR3-Go
aa
3,165 movies
Motion Correction (RELION 4.0)
Patch CTF estimation
3,165 micrographs
1,384,864 particles
2D classification
Heterogeneous refinement
Re-extraction
363,327 particles
Heterogeneous refinement
Non-uniform refinement
3.14 Å 221,220 particles
3D classification without alignment (RELION 4.0)
Bayesian polishing
CTF refinement
Non-uniform refinement
Sharpening
(Phenix.auto_sharpen)
3.03 Å 116,462 particles
Local refinement
focused on receptor
Sharpening
(Phenix.auto_sharpen)
3.16 Å 116,462 particles
ab
500Å
ac
ad
ae
GSFSC Resolution: 3.03Å
af
ag
ah
GSFSC Resolution: 3.16Å
ai
ba
3,030 movies
Motion Correction (RELION 4.0)
Patch CTF estimation
3,030 micrographs
1,527,953 particles
2D classification
Heterogeneous refinement
Re-extraction
360,233 particles
Heterogeneous refinement
Non-uniform refinement
3.23 Å 182,526 particles
3D classification without alignment (RELION 4.0)
Bayesian polishing
CTF refinement
Non-uniform refinement
Sharpening
(Phenix.auto_sharpen)
3.07 Å 150,213 particles
Local refinement
focused on receptor
Sharpening
(Phenix.auto_sharpen)
3.53 Å 150,213 particles
bb
500Å
bc
bd
be
GSFSC Resolution: 3.07Å
bf
bg
bh
GSFSC Resolution: 3.53Å
bi
ca
3,125 movies
Motion Correction (RELION 4.0)
Patch CTF estimation
3,125 micrographs
1,633,141 particles
2D classification
Heterogeneous refinement
Re-extraction
298,771 particles
Heterogeneous refinement
Non-uniform refinement
3.24 Å 173,083 particles
3D classification without alignment (RELION 4.0)
Bayesian polishing
CTF refinement
Non-uniform refinement
Sharpening
(Phenix.auto_sharpen)
3.30 Å 41,722 particles
Local refinement
focused on receptor
Sharpening
(Phenix.auto_sharpen)
3.68 Å 41,722 particles
cb
500Å
cc
cd
ce
GSFSC Resolution: 3.30Å
cf
cg
ch
GSFSC Resolution: 3.68Å
ci
90°

**Extended Data Fig. 10: Cryo-EM data processing pipeline of VUF10661-CXCR3-Go, VUF11418-CXCR3-Go and apo-CXCR3-Go.**

**aa, ba, ca,** Schematic representations of the cryo-EM data processing workflow for VUF10661-CXCR3-Go (**aa**), VUF11418-CXCR3-Go (**ba**) and apo-CXCR3-Go (**ca**).

**ab, bb, cb,** Representative cryo-EM images of the VUF10661–CXCR3–Go (**ab**), VUF11418–CXCR3–Go (**bb**) and apo-CXCR3-Go (**cb**) recorded on a 300 kV Titan Krios with a K3 camera.

**ac, bc, cc,** Representative 2D averages of curated particles for VUF10661-CXCR3-Go (**ac**), VUF11418-CXCR3-Go (**bc**) and apo-CXCR3-Go (**cc**).

**ad, bd, cd,** Local resolution of the overall refined maps for VUF10661-CXCR3-Go (**ad**), VUF11418-CXCR3-Go (**bd**) and apo-CXCR3-Go (**cd**).

**ae, be, ce,** Gold standard fourier shell correlation curve (FSC) at 0.143 threshold for overall refined maps of VUF10661-CXCR3-Go (**ae**), VUF11418-CXCR3-Go (**be**) and apo-CXCR3-Go (**ce**).

**af, bf, cf,** Angular distribution of the overall refinement for VUF10661-CXCR3-Go (**af**), VUF11418-CXCR3-Go (**bf**) and apo-CXCR3-Go (**cf**).

**ag, bg, cg,** Local resolution of the local refined maps for VUF10661-CXCR3-Go (**ag**), VUF11418-CXCR3-Go (**bg**) and apo-CXCR3-Go (**cg**).

**ah, bh, ch,** Gold standard fourier shell correlation curve (FSC) at 0.143 threshold for local refined maps of VUF10661-CXCR3-Go (**ah**), VUF11418-CXCR3-Go (**bh**) and apo-CXCR3-Go (**ch**).

**ai, bi, ci,** Angular distribution of the local refinement for VUF10661-CXCR3-Go (**ai**), VUF11418-CXCR3-Go (**bi**) and apo-CXCR3-Go (**ci**).

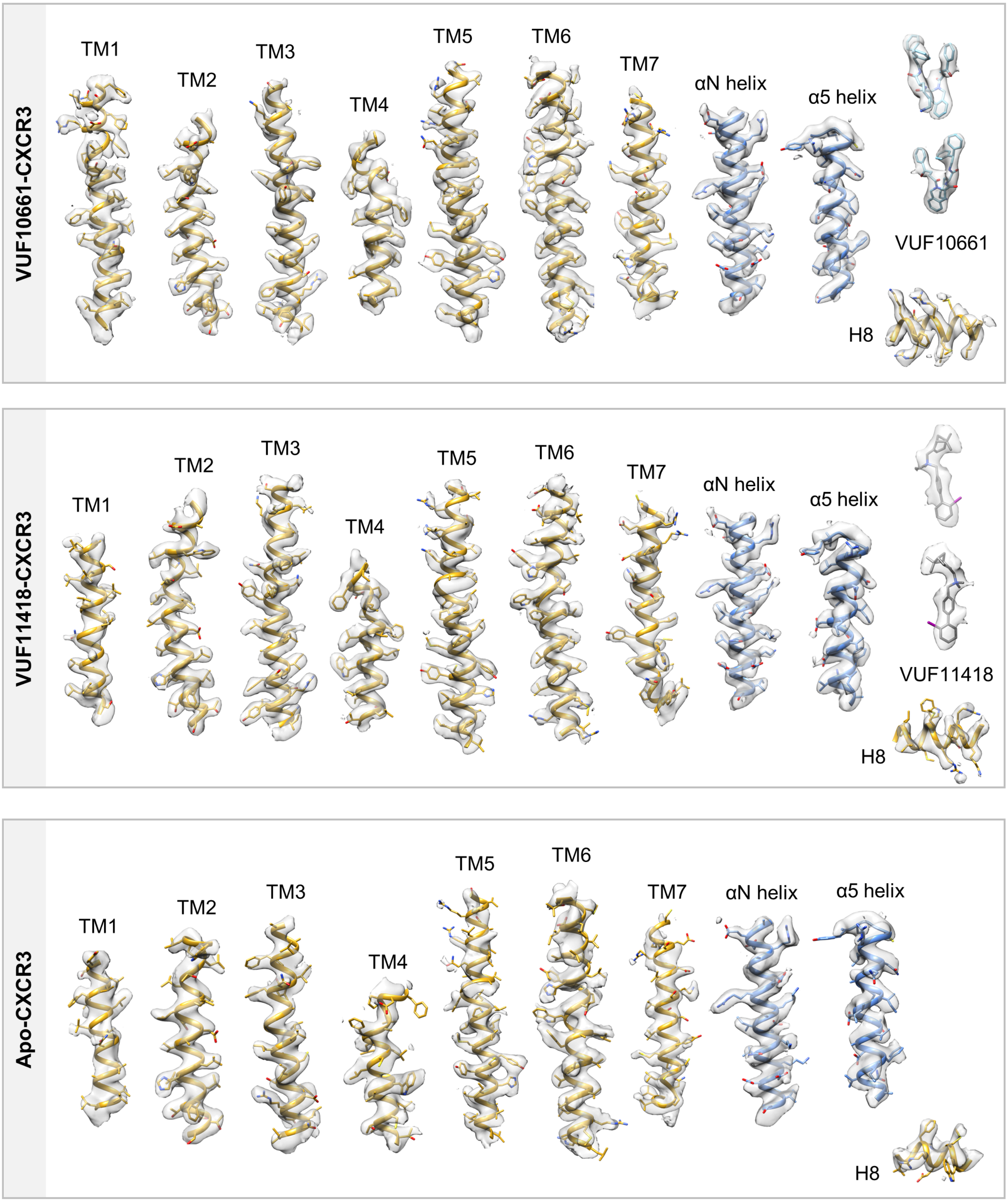


**Extended Data Fig.11: Exemplary electron density maps of the CXCR3 complexes.**

EM densities of the TMs, Helix 8, αN helix and α5 helix of VUF10661-CXCR3, VUF11418-CXCR3 and Apo-CXCR3, and VUF10661 and VUF11418.

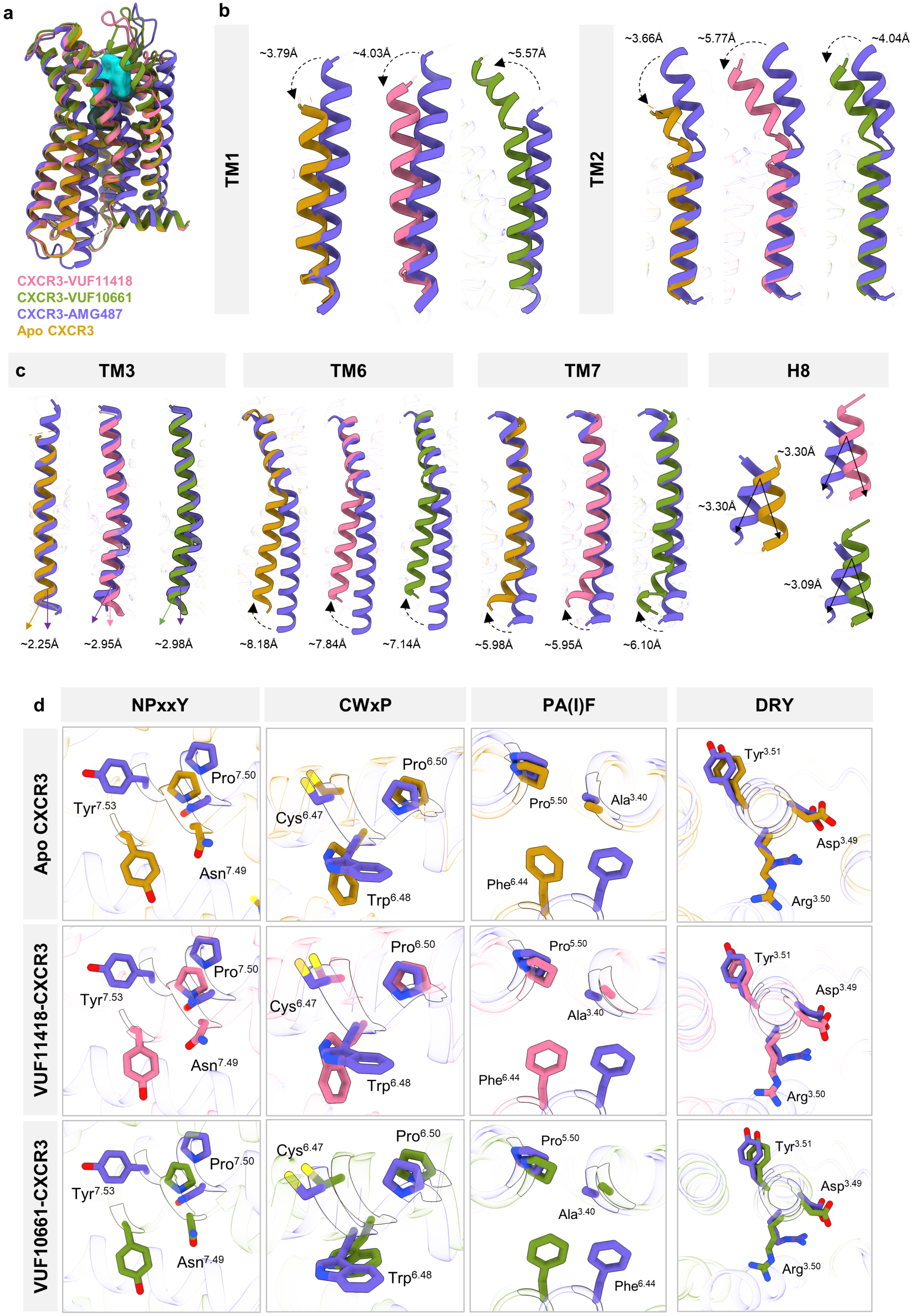
a
CXCR3-VUF11418
CXCR3-VUF10661
CXCR3-AMG487
Apo CXCR3
b
TM1
~3.79Å
~4.03Å
~5.57Å
TM2
~3.66Å
~5.77Å
~4.04Å
c
TM3
TM6
TM7
H8
~2.25Å
~2.95Å
~2.98Å
~8.18Å
~7.84Å
~7.14Å
~5.98Å
~5.95Å
~6.10Å
~3.30Å
~3.30Å
~3.09Å
d
NPxxY
CWxP
PA(I)F
DRY
Apo CXCR3
VUF11418-CXCR3
VUF10661-CXCR3
Tyr7.53
Pro7.50
Asn7.49
Cys6.47
Pro6.50
Trp6.48
Pro5.50
Ala3.40
Phe6.44
Tyr3.51
Asp3.49
Arg3.50

**Extended Data Fig. 12: Major conformational changes on CXCR3 activation.**

**a,** Superimposition of inactive CXCR3 with apo CXCR3 receptor and CXCR3 receptor bound to VUF11418 and VUF10661 respectively. **b, c,** Displacements of TM1, TM3, TM6, TM7 and helix8 upon CXCR3 activation in the structures of apo-CXCR3, VUF11418-CXCR3, VUF10661-CXCR3 respectively. **d,** Conformational changes in the conserved microswitches (DRY, PIF, NPxxY, CWxP) in the active structure of CXCR3.

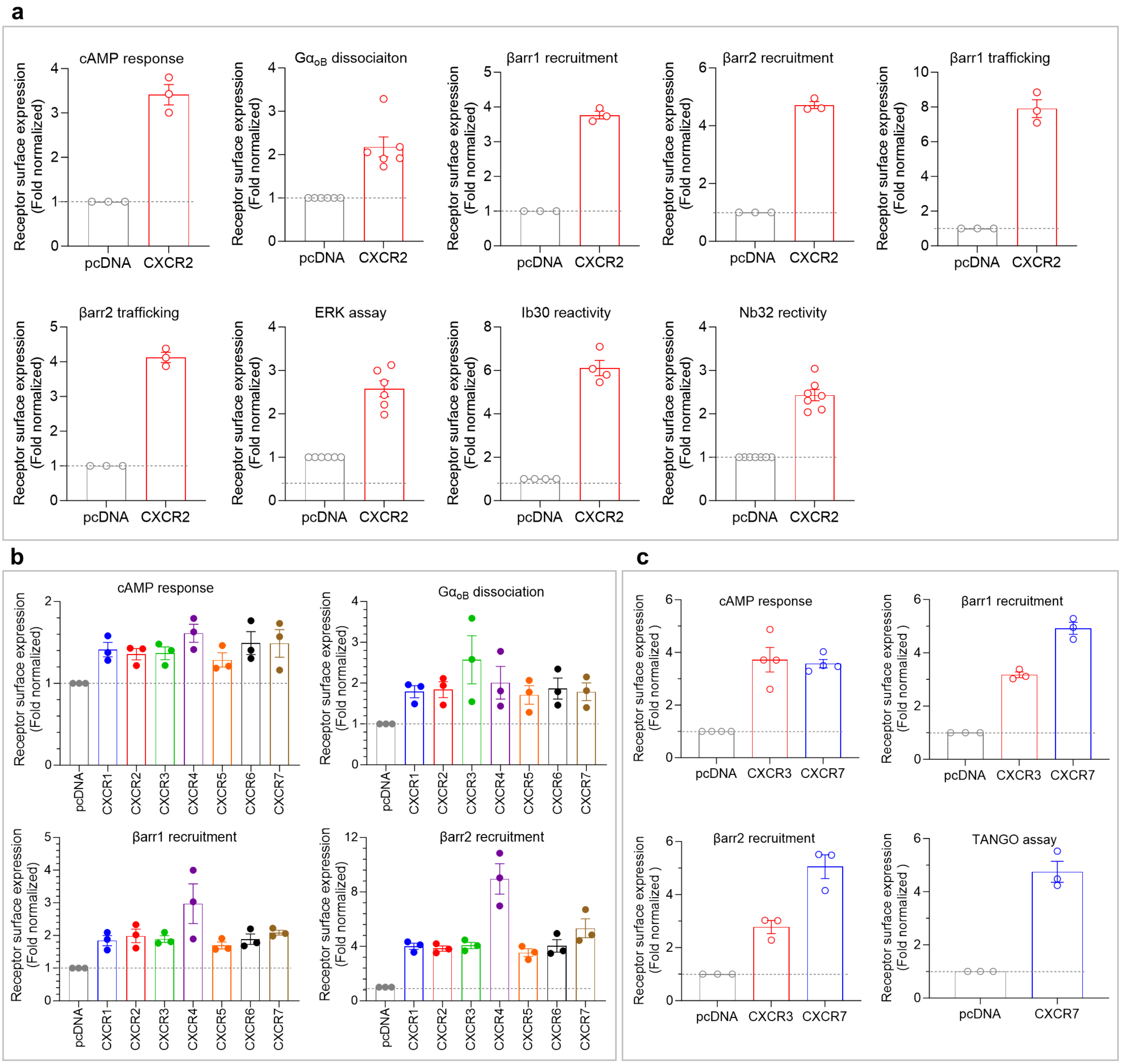


**Extended Data Fig. 13: Surface expression of receptors in various assays.**

**a,** Surface expression of CXCR2 in various assays performed. Data (mean±SEM) represents three to six independent biological replicates. **b,** All the receptors used in screening VUF10661 and VUF11418 showed robust expression. Data (mean±SEM) represents three independent biological replicates. **c,** CXCR3 and CXCR7 showed comparable levels of surface expression. Data (mean±SEM) represents three to four independent biological replicates.
